# Metagenomic Characteristics of Bacterial Response to Petroleum Hydrocarbon Contamination in Diverse Environments as Revealed by Functional Taxonomic Strategies

**DOI:** 10.1101/097063

**Authors:** Arghya Mukherjee, Bobby Chettri, James S. Langpoklakpam, Pijush Basak, Aravind Prasad, Ashis K. Mukherjee, Maitree Bhattacharyya, Arvind K. Singh, Dhrubajyoti Chattopadhyay

## Abstract

Microbial remediation of oil polluted habitats remains one of the foremost methods for restoration of petroleum hydrocarbon contaminated environments. The development of effective bioremediation strategies however, require an extensive understanding of the resident microbiome of these habitats. Recent developments such as high-throughput sequencing has greatly facilitated the advancement of microbial ecological studies in oil polluted habitats. However, effective interpretation of biological characteristics from these large datasets remains a considerable challenge. In this study, we have implemented recently developed bioinformatic tools for analyzing 65 publicly available 16S rRNA datasets from 12 diverse hydrocarbon polluted habitats to decipher metagenomic characteristics of bacterial communities of the same. We have comprehensively described phylogenetic and functional compositions of these habitats and additionally inferred a multitude of metagenomic features including 255 taxa and 414 functional modules which can be used as biomarkers for effective distinction between the 12 oil polluted sites. We have identified essential metabolic signatures and also showed that significantly over-represented taxa often contribute to either or both, hydrocarbon degradation and additional important functions. Our findings reveal significant differences between hydrocarbon contaminated sites and establishes the importance of endemic factors in addition to petroleum hydrocarbons as driving factors for sculpting hydrocarbon contaminated bacteriomes.

## Introduction

Anthropogenic activities and agents leading to contamination of the environment is one of the major issues that developing and developed industrial societies face today. Petroleum hydrocarbons are the most widespread of these anthropogenic agents and frequently contaminate aquatic and terrestrial ecosystems through releases of hydrocarbon during production, operational use, and transportation. The development, effectiveness and availability of technologies and strategies pose a significant challenge for the remediation, rehabilitation and restoration of these contaminated environments. Many of the technologies developed and in use for the restoration of oil contaminated environments exploit the potential of biological systems, in particular microbial systems, to use these toxic compounds as substrates for growth. Hence, much of the research conducted on bioremediation has concentrated on the capabilities of a single or couple of microbes exhibiting robust and effective growth using petroleum hydrocarbons. However, in the environment, bioremediation is often a complex process involving co-metabolism, cross-induction, inhibition and non-interaction among microbes ^1-3^, possibly as petroleum hydrocarbons are a mixture of organic pollutants and therefore are used differently by different microbes. These findings, along with others, established bioremediation as a process mediated by a consortium of microbes rather than a few. Thus, characterization of microbial communities of oil contaminated environments could potentially provide guidelines for effective remediation and restoration of such environments.

Until recently, it was only possible to study a handful of microorganisms of interest isolated from source materials (as blood, soil, water or air), given the restrictions of the composition of culture media which cannot reflect and mimic the dynamic nutrient fluxes of the source environment. Indeed, only 1% of microorganisms were found to be cultivable using a set of media from the highly characterized soil rhizosphere ^4^. The advent of high throughput massively-parallel sequencing methods has however, allowed us to investigate the entire complement of organisms inhabiting a certain environment. These next-generation sequencing methods (NGS) include a variety of methods to holistically study any biological system such as amplicon sequencing (for variant identification and phylogenetic surveys), whole genome shotgun sequencing (single organism genome and metagenomes) and RNA-Seq (transcriptomes, metatranscriptomes and identification of non-regular RNAs). These powerful methods have ushered in rapid advances in bioinformatics approaches leading to development of software capable of handling huge amounts of data and offering meaningful biological interpretations of the same. Although a technological breakthrough in modern science, a number of NGS methods as metagenomic and transcriptomic/metatranscriptomic sequencing are still expensive and hence, most studies on ecological processes on bioremediation report marker surveys as 16S rRNA gene amplicon sequencing when dealing with a large number of samples. Thus, in general, most of these studies concentrated on interpretations from microbial community composition but inferred poorly regarding functional and metabolic properties of the same.

Recently, with the implementation of the Human Microbiome Project (HMP), bioinformatic advancements have been furthered through the development of powerful new computational tools for effective interpretation and visualization of taxonomic and functional composition of microbial communities ^5,6^ These tools have obvious applications for the analysis of huge amounts of microbial genomic/amplicon/transcriptomic data collected from other sources such as soil, water and so on. Some particularly interesting computational tools allow to explain the complex mutual interactions and heterogeneit inherent in microbial communities through network-based correlation analyses ^7^, prediction of metagenomic biomarkers ^8^ and prediction of metagenomes from 16S rRNA data ^9^.

It is well understood that depending on the environment, the method of bioremediation will vary. However, essential information required for development of these technologies include the response of microbes to petroleum hydrocarbons and their dynamics with the immediate environment. Unfortunately, despite the large amount of work done on microbial community composition across a myriad of oil contaminated environments, mainly through 16S rRNA amplicon sequencing, no attempt has been made to find differential metagenomic signatures among these studies. In the present study, we have aimed to investigate the taxonomic and functional characteristics of diverse oil contaminated environments using recent bioinformatics tools through an evolving pipeline to process metagenomic data. Due to the current paucity of metagenomic datasets for this kind of study and for the large availability of them, we used 61 publicly available 16S rRNA datasets and 4 from this study as inputs for our analysis. Consequently, metagenomic level characteristics of bacterial composition and metabolic potential were comprehensively deduced for oil contaminated soils in north-east India along with 11 other petroleum hydrocarbon contaminated habitats. We inferred an array of differentially abundant taxonomic and functional features which may be used as biomarkers for successful distinction of different oil contaminated habitats as well as for monitoring of bioremediation efforts in the same. Additionally, we deduced important metabolic pathways for all contaminated environments. Evaluation of correlation between taxa and functional orthologs was also carried out along with estimation of metagenomic contributions to hydrocarbon degradation to detect taxa responsible for critical functions in oil polluted habitats. Furthermore, a network of bacterial interaction patterns was inferred to deduce complex co-occurrence and co-exclusion relationships in these environments. We found that phylogenetic and functional composition oil contaminated bacteriomes were significantly different to each other and greatly influenced by immediate environmental factors along with petroleum hydrocarbon contamination. Our investigation provides novel and valuable insights into the differential nature of various oil polluted habitats and hopefully improves upon previous understanding of these environments.

## Materials and Methods

### Ethics statement

No specific permits were required for the samplings carried out at the fields described in Noonmati or Barhola in Assam, India. The study sites are not privately owned or protected in any terms. Also, the field work did not involve any protected or endangered species.

### Collection of soil samples

Oil contaminated soil samples were collected from Noonmati Oil Refinery in Guwahati and from oilfields in Barhola, both in Assam, India. Soil samples were collected in triplicate from both sites from the surface (0-10 cm) and beneath (20-40 cm) using sterile equipment. The soil samples were transported to the laboratory on ice within 48-72 hours. In the laboratory, replicates for each sample was mixed and homogenized to uniformity to form four composite samples prior to isolation of soil DNA.

### Extraction of total soil DNA

Soil DNA was isolated using the PowerMax Soil DNA Isolation Kit (MoBio Labs, Carlsbad, CA, USA) according to the manufacturer's protocol. Integrity of isolated soil DNA was then checked in a 0.8% agarose gel, while a NanoDrop 2000 Spectrophotometer (Thermo Scientific, Wilmington, DE, USA) was used to obtain qualitative (260:280 and 260:230 ratios) and quantitative estimates.

### 16S rRNA PCR Amplification and Pyrosequencing

The V1-V3 region of the bacterial 16S rRNA gene was amplified using specially designed fusion primers, consisting of both template derived sequence and a specific sequence as recommended by the “Sequencing Technical Bulletin No. 013-2009” (454 Life Sciences, Branford, CT, USA). The Fusion forward primer (5’-3’) consisted of a Roche A adapter (CCATCTCATCCCTGCGTGTCTCCGAC), a key sequence (TCAG), a 10 bp Multiplex Identifier (MID) sequence and a template specific sequence (GAGTTTGATCMTGGCTCAG) derived from the universal 16S rRNA primer 27F ^10^. The Fusion reverse primer consisted of a Roche B adapter (CCTATCCCCTGTGTGCCTTGGCAGTC), a key sequence (TCAG) and a template specific sequence (ATTACCGCGGCTGCTGG) derived from the universal 16S rRNA gene primer 541R ^11^. V1-V3 region of the 16S rRNA gene was amplified through PCR in a 25 μl reaction mix containing 2.5 μl Fast Start Buffer (10X), 1 μl each of Fusion forward primer (10 μM) and Fusion reverse primer (10 μM), 0.5 μl dNTPs (10 mM), 0.5 μl Fast Start Taq Polymerase (5U/μl Fast Start High Fidelity PCR System, Roche) with the rest made up with water. The PCR was carried out in a Veriti Thermal Cycler (Thermo Scientific, Wilmington, DE, USA) under the following conditions: initial denaturation at 94°C for 3 minutes, followed by 25 cycles of denaturation at 94 °C for 15 secs, annealing at 58°C for 45 secs and elongation at 72 °C for 1 min, with a final elongation at 72°C for 10 mins. Thereafter samples were stored at -20 °C, if required.

### Sequence processing and taxonomic analysis of 16S rRNA data

Raw 16S rRNA sequencing reads were checked for quality using FastQC ^12^ and subsequently processed using mothur ^13^, which included trimming of adapters, keys, MIDs and primers from the raw sequences. Sequences were further filtered for quality using the following non-default parameters: maxhomop = 6, maxambig = 0, maxlength = 575, minlength = 200, qwindowaverage = 30, bdiffs = 1, pdiffs = 2, and tdiffs = 2. Filtered high quality sequences were then aligned to the mothur implementation of the SILVA database and trimmed for the alignment region. Chimeric sequences were then removed from the datasets using the mothur implementation of Uchime ^14^. Filtered sequences were then taxonomically classified using the May 2013 release of the Greengenes database ^15^ and contaminating archaeal, eukaryal, mitochondrial and chloroplast sequences or sequences classified as unknown were removed from further analysis. mothur was further used to compute coverage, boneh index (for additional 500 sequences), observed species richness, and alpha diversity metrics through the estimation OTUs at 0.03 level of phylogenetic divergence. OTUs were further classified using the taxonomy file generated in the steps before. Taxonomically classified OTUs were converted to number of sequences and visualized as circular cladograms using the standalone graphical tool GraPhlan v0.95 ^16^.

### Collection and quality filtering of 16S rRNA datasets from oil contaminated environments

Sixty-one 16S rRNA datasets on oil degradation studies from 11 different environments collected from publicly available resources along with four samples from this study were used for the present study (Table 1, Supplementary Table S1). These included four datasets representing upper soil layers of the Tundra biome (Tu), four from subsurface layers of the Tundra biome (Tb), four from the permafrost layers of the Tundra (Tp), nine from surface soil of Chinese oil refineries (C), twelve representing different regions of the arctic biome (A), four from surface soils of Indian oil refineries (I), three from mangroves (M), seven from surficial marine sediments (DWH), seven from oil sands cores (OSC), four from surface waters of oil sands tailings ponds (OSTPu), three from oil sands tailings pond waters at median depth (OSTPm) and four from deep oil sands tailings pond waters (OSTPd). We deliberately kept the taiga and OSTP samples separate even though we expected high amounts of similarity between them in certain aspects when compared to other samples, due to evidence of ample distinctive characteristics in the said samples in their parent studies ^17,18^. All the 16S rRNA datasets used can be downloaded through the list of accession numbers provided in Supplementary Table S1. All datasets used in the study presented, were sequenced in either Roche 454, Illumina or ABI Ion Torrent platforms. The 16S rRNA datasets are described in greater detail in Table 1. The downloaded 16S rRNA datasets were checked for quality using FastQC and filtered for high quality sequences in mothur using the following criteria: minimum sequence length of 100 bp, sequences trimmed when average quality drops below 20 in a sliding window of 15 bp, and a maximum of 2 mismatches in the barcode-key-template region of the reads.

**Table 1.** Summary of datasets used in the study. (For additional details, refer to Supplementary data Table S1).

| Biome Type | ID | Sequencing Platform | Original ID | Location | Depth of sample collection (cm below surface) | Source material for sequencing | Predominant contaminant/hydrocarbon | Reference |
| --- | --- | --- | --- | --- | --- | --- | --- | --- |
| Urban | I1 | 454 GS Junior | Noonmati_Surface | Guwahati, Assam, India | 0-10 | *In situ* soil | Crude oil | This study |
| Urban | I2 | 455 GS Junior | Noonmati_Deep | Guwahati, Assam, India | 20-30 | *In situ* soil | Crude oil | This study |
| Urban | I3 | 456 GS Junior | Barhola_Surface | Barhola, Assam, India | 0-10 | *In situ* soil | Crude oil | This study |
| Urban | I4 | 457 GS Junior | Barhola_Deep | Barhola, Assam, India | 20-30 | *In situ* soil | Crude oil | This study |
| Arctic | A1 | Ion Torrent PGM | AH1d1, AH1d2 | Axel Heiburg, Nunavut, Canada | 0-15 | Treated microcosm sediment | Diesel oil | <sup>78</sup> |
| Arctic | A2 | Ion Torrent PGM | AK1d1, AK1d2, AK1d3 | Toolik Lake, Alaska, USA | 0-15 | Treated microcosm sediment | Diesel oil | <sup>78</sup> |
| Arctic | A3 | Ion Torrent<br>PGM | AL1d1, AL1d2,<br>AL1d3 | Alert,<br>Nunavut,<br>Canada | 0-15 | Treated<br>microcosm<br>sediment | Diesel oil | 78 |
| Arctic | A4 | Ion Torrent<br>PGM | AVKd1, AVKd2,<br>AVKd3 | Akulivik,<br>Quebec,<br>Canada | 0-15 | Treated<br>microcosm<br>sediment | Diesel oil | 78 |
| Arctic | A5 | Ion Torrent<br>PGM | BDEd1, BDEd2,<br>BDEd3 | Baie<br>Deception,<br>Quebec,<br>Canada | 0-15 | Treated<br>microcosm<br>sediment | Diesel oil | 78 |
| Arctic | A6 | Ion Torrent<br>PGM | BY1d1, BY1d2,<br>BY1d3 | Bylot Island,<br>Nunavut,<br>Canada | 0-15 | Treated<br>microcosm<br>sediment | Diesel oil | 78 |
| Arctic | A7 | Ion Torrent<br>PGM | EBAd1, EBAd2,<br>EBAd3 | East Bay,<br>Nunavut,<br>Canada | 0-15 | Treated<br>microcosm<br>sediment | Diesel oil | 78 |
| Arctic | A8 | Ion Torrent<br>PGM | IQAd1, IQAd2,<br>IQAd3 | Iqaluit,<br>Nunavut,<br>Canada | 0-15 | Treated<br>microcosm<br>sediment | Diesel oil | 78 |
| Arctic | A9 | Ion Torrent<br>PGM | NORd1, NORd2,<br>NORd3 | Tromso,<br>Norway | 0-15 | Treated<br>microcosm<br>sediment | Diesel oil | 78 |
| Arctic | A10 | Ion Torrent<br>PGM | RANd1, RANd2,<br>RANd3 | Rankin<br>Inlet,<br>Nunavut,<br>Canada | 0-15 | Treated<br>microcosm<br>sediment | Diesel oil | 78 |
| Arctic | A11 | Ion Torrent<br>PGM | RUSd1, RUSd2,<br>RUSd3 | Yamal,<br>Russia | 0-15 | Treated<br>microcosm<br>sediment | Diesel oil | 78 |
| Arctic | A12 | Ion Torrent PGM | THUd1, THUd2, THUd3 | Thule, Greenland | 0-15 | Treated microcosm sediment | Diesel oil | 78 |
| Urban | C1 | Illumina Miseq | CQM1 | Changqing, China | 2-10 | *In situ* soil | Crude oil | 79 |
| Urban | C2 | Illumina Miseq | CQM2 | Changqing, China | 2-10 | *In situ* soil | Crude oil | 79 |
| Urban | C3 | Illumina Miseq | CQM3 | Changqing, China | 2-10 | *In situ* soil | Crude oil | 79 |
| Urban | C4 | Illumina Miseq | DQM3 | Daqing, China | 2-10 | *In situ* soil | Crude oil | 79 |
| Urban | C5 | Illumina Miseq | DQM4 | Daqing, China | 2-10 | *In situ* soil | Crude oil | 79 |
| Urban | C6 | Illumina Miseq | DQM5 | Daqing, China | 2-10 | *In situ* soil | Crude oil | 79 |
| Urban | C7 | Illumina Miseq | DQM12 | Daqing, China | 2-10 | *In situ* soil | Crude oil | 79 |
| Urban | C8 | Illumina Miseq | DQM50 | Daqing, China | 2-10 | *In situ* soil | Crude oil | 79 |
| Urban | C9 | Illumina Miseq | DQM200 | Daqing, China | 2-10 | *In situ* soil | Crude oil | 79 |
| Mangrove | M1 | 454 GS FLX | T23 2% I, T23 2% II | Restinga da Marambaia, Rio de Janeiro, Brazil | 0-20 | Treated microcosm sediment | Crude oil | 43 |
| Mangrove | M2 | 455 GS FLX | T66 2% I, T66 2% II | Restinga da Marambaia, Rio de | 0-20 | Treated microcosm sediment | Crude oil | 43 |
|  |  |  |  | Janeiro,<br>Brazil |  |  |  |  |
| Mangrove | M3 | 456 GS FLX | T23 5% I, T23 5% II | Restinga da<br>Marambaia,<br>Rio de<br>Janeiro,<br>Brazil | 0-20 | Treated<br>microcosm<br>sediment | Crude oil | 43 |
| Marine<br>sediment | DWH1 | Illumina | SE-20101001-GY-LBNL1-BC-120 | Gulf of<br>Mexico | 0-1<sup>#</sup> | *In situ* soil | Crude oil | 47 |
| Marine<br>sediment | DWH2 | Illumina | SE-20101001-GY-ALTNF001-BC-139 | Gulf of<br>Mexico | 0-1<sup>#</sup> | *In situ* soil | Crude oil | 47 |
| Marine<br>sediment | DWH3 | Illumina | SE-20101001-GY-NF006MOD-BC-143 | Gulf of<br>Mexico | 0-1<sup>#</sup> | *In situ* soil | Crude oil | 47 |
| Marine<br>sediment | DWH4 | Illumina | SE-20101017-GY-D031S-BC-278 | Gulf of<br>Mexico | 0-1<sup>#</sup> | *In situ* soil | Crude oil | 47 |
| Marine<br>sediment | DWH5 | Illumina | SE-20101017-GY-D040S-BC-315 | Gulf of<br>Mexico | 0-1<sup>#</sup> | *In situ* soil | Crude oil | 47 |
| Marine<br>sediment | DWH6 | Illumina | SE-20101017-GY-D038SW-BC-331 | Gulf of<br>Mexico | 0-1<sup>#</sup> | *In situ* soil | Crude oil | 47 |
| Marine<br>sediment | DWH7 | Illumina | SE-20101017-GY-D042S-BC-350 | Gulf of<br>Mexico | 0-1<sup>#</sup> | *In situ* soil | Crude oil | 47 |
| Taiga | Tu1 | 456 GS FLX | 1330 | Walagan,<br>China | 20-30 | Treated<br>microcosm<br>sediment | Crude oil | 18 |
| Taiga | Tu2 | 456 GS FLX | 2330 | Walagan<br>North,<br>China | 20-30 | Treated<br>microcosm<br>sediment | Crude oil | 18 |
| Taiga | Tu3 | 456 GS FLX | 3330 | Taiyuan,<br>China | 20-30 | Treated<br>microcosm<br>sediment | Crude oil | 18 |
| Taiga | Tu4 | 456 GS FLX | 4330 | Jiagedaqi,<br>China | 20-30 | Treated<br>microcosm<br>sediment | Crude oil | 18 |
| Taiga | Tb1 | 456 GS FLX | 1530 | Walagan,<br>China | 70-80 | Treated<br>microcosm<br>sediment | Crude oil | 18 |
| Taiga | Tb2 | 456 GS FLX | 2530 | Walagan<br>North,<br>China | 70-80 | Treated<br>microcosm<br>sediment | Crude oil | 18 |
| Taiga | Tb3 | 456 GS FLX | 3530 | Taiyuan,<br>China | 70-80 | Treated<br>microcosm<br>sediment | Crude oil | 18 |
| Taiga | Tb4 | 456 GS FLX | 4530 | Jiagedaqi,<br>China | 70-80 | Treated<br>microcosm<br>sediment | Crude oil | 18 |
| Taiga | Tp1 | 456 GS FLX | 1630 | Walagan,<br>China | 140-150 | Treated<br>microcosm<br>sediment | Crude oil | 18 |
| Taiga | Tp2 | 456 GS FLX | 2630 | Walagan<br>North,<br>China | 140-150 | Treated<br>microcosm<br>sediment | Crude oil | 18 |
| Taiga | Tp3 | 456 GS FLX | 3630 | Taiyuan,<br>China | 140-150 | Treated<br>microcosm<br>sediment | Crude oil | 18 |
| Taiga | Tp4 | 456 GS FLX | 4630 | Jiagedaqi,<br>China | 140-150 | Treated<br>microcosm<br>sediment | Crude oil | 18 |
| Arctic | OSC1 | 456 GS FLX | 2010Suncor Oil Sands Core Run 11 Subsample 1 | Alberta, Canada | 2,985-2,990 | *In situ* soil | Oil sands bitumen | 17 |
| Arctic | OSC3 | 456 GS FLX | 2010Suncor Oil Sands Core Run 11 Subsample 3 | Alberta, Canada | 2,985-2,990 | *In situ* soil | Oil sands bitumen | 17 |
| Arctic | OSC4 | 456 GS FLX | 2010Suncor Oil Sands Core Run 11 Subsample 4 | Alberta, Canada | 2,985-2,990 | *In situ* soil | Oil sands bitumen | 17 |
| Arctic | OSC5 | 456 GS FLX | 2010Suncor Oil Sands Core Run 11 Subsample 5 | Alberta, Canada | 2,985-2,990 | *In situ* soil | Oil sands bitumen | 17 |
| Arctic | OSC7 | 456 GS FLX | 2010Suncor Oil Sands Core Run 11 Subsample 7 | Alberta, Canada | 2,985-2,990 | *In situ* soil | Oil sands bitumen | 17 |
| Arctic | OSC9 | 456 GS FLX | 2010Suncor Oil Sands Core Run 11 Subsample 9 | Alberta, Canada | 2,985-2,990 | *In situ* soil | Oil sands bitumen | 17 |
| Arctic | OSC12 | 456 GS FLX | 2010Suncor Oil Sands Core Run 11 Subsample 13 | Alberta, Canada | 2,985-2,990 | *In situ* soil | Oil sands bitumen | 17 |
| Arctic | OSTPu1 | 456 GS FLX | 2009Suncor Tailings Pond 5 (6.5 ft) | Alberta, Canada | 200 | *In situ* soil | Oil sands bitumen | 17 |
| Arctic | OSTPu2 | 456 GS FLX | 2009Suncor Tailings Pond 5 (8.0 ft) | Alberta, Canada | 240 | *In situ* soil | Bitumen and various other hydrocarbons | 17 |
| Arctic | OSTPu3 | 456 GS FLX | 2010Syncrude Tailings Pond MLSB M MFT - 1m | Alberta, Canada | 100 | *In situ* soil | Bitumen and various other hydrocarbons | 17 |
| Arctic | OSTPu4 | 456 GS FLX | 2010Syncrude<br>Tailings Pond MLSB<br>N MFT - 1.1m | Alberta,<br>Canada | 110 | *In situ* soil | Bitumen and various other<br>hydrocarbons | 17 |
| Arctic | OSTPm2 | 456 GS FLX | 2009Suncor<br>Tailings Pond 5 (25<br>ft) | Alberta,<br>Canada | 750 | *In situ* soil | Bitumen and various other<br>hydrocarbons | 17 |
| Arctic | OSTPm4 | 456 GS FLX | 011Suncor Tailings<br>Pond 6 (7 m) | Alberta,<br>Canada | 700 | *In situ* soil | Bitumen and various other<br>hydrocarbons | 17 |
| Arctic | OSTPm6 | 456 GS FLX | 2010Syncrude<br>Tailings Pond MLSB<br>N MFT - 6.1m | Alberta,<br>Canada | 610 | *In situ* soil | Bitumen and various other<br>hydrocarbons | 17 |
| Arctic | OSTPd1 | 456 GS FLX | 2009Suncor<br>Tailings Pond 5 (40<br>ft) | Alberta,<br>Canada | 1220 | *In situ* soil | Bitumen and various other<br>hydrocarbons | 17 |
| Arctic | OSTPd2 | 456 GS FLX | 2009SuncorTailings<br>Pond 5 (45 ft) | Alberta,<br>Canada | 1370 | *In situ* soil | Bitumen and various other<br>hydrocarbons | 17 |
| Arctic | OSTPd3 | 456 GS FLX | 2010Suncor<br>Tailings Pond 6 (12<br>m) | Alberta,<br>Canada | 1200 | *In situ* soil | Bitumen and various other<br>hydrocarbons | 17 |
| Arctic | OSTPd4 | 456 GS FLX | 2011Suncor<br>Tailings Pond 6 (13<br>m) | Alberta,<br>Canada | 1300 | *In situ* soil | Bitumen and various other<br>hydrocarbons | 17 |

#All samples collected at an average of ~1500 metres below sea level, depth given is from surface of ocean floor marine sediment.

### Analysis of microbial community structure and composition in 16S rRNA datasets

mothur was used to estimate abundances of bacterial taxa in the 16S rRNA datasets as described above. Briefly, all the datasets containing high quality reads were aligned against the mothur implementation of the SILVA 16S rRNA database, followed by removal of chimeric sequences using the mothur implementation of Uchime. Classification of the filtered sequences against the Greengenes database was carried out then, upon which contaminating archaeal, chloroplast, mitochondrial, eukaryal or unknown sequences were removed. Finally, OTUs were predicted from these high quality sequences. OTUs were again mapped to the sequence taxonomy file generated previously in mothur to generate comparative taxonomy data for the datasets. We also assessed the compositional similarity between the soil samples from different sites. For doing this, we compared the pairwise taxonomic abundances from each site against each other and within the datasets as well, using Bray-Curtis measure for estimation of beta diversity ^19^. The permutation-based multivariate analysis of variance (PERMANOVA) was used to test the homogeneity of taxonomic dispersion across samples along with concomitant estimation of 2D stress. The resulting Bray-Curtis similarity distance matrix was used as input for ordination of the oil contaminated samples through non-metric multidimensional scaling (NMDS) in PAST v3.11 ^20^.

### Metagenome prediction and metabolic reconstruction of 16S rRNA datasets

Metagenomes were predicted from 16S rRNA data using PICRUSt ^9^. OTU data generated in mothur for all 16S rRNA datasets was used to prepare .*biom* files formatted as input for PICRUSt v1.1.0 ^9^ with the *make.biom* script available in mothur. PICRUSt requires OTU abundances mapped to Greengenes OTU IDs as input for prediction of corresponding metagenomes. PICRUSt databases for 16S rRNA copy number normalization and KEGG ortholog prediction were updated using publicly available information listed in Integrated Microbial Genomes (IMG)^21^ as on 4^th^ April, 2016, according to the instructions (default settings) provided in the Genome Prediction Tutorial for PICRUSt (http://picrust.github.io/picrust/tutorials/genome_prediction.html#genome-prediction-tutorial). The update involved the inclusion of 16S rRNA copy number information and KEGG ortholog (KO) annotation data as per KEGG v77.1 ^22^ for ~34,000 bacterial and archaeal genomes available in IMG. 16S rRNA copy numbers for 16S rRNA datasets were normalized using the *normalize*_*by*_*copy*_*number*.*py* script. Metagenomes were predicted from the copy number normalized 16S rRNA data in PICRUSt using the *predict*_*metagenomes*.*py* script against the updated and PICRUSt-formatted, characterized protein functional database of KEGG Orthology. Contributions of various taxa to different KOs were computed with the script *metagenome*_*contributions*.*py* and visualized with the script *plot*_*metagenome*_*contributions*.*R* (https://groups.google.com/forum/#!topic/picrust-users/Hq9_G23J9W4) and ggplot ^23^ in R (http://www.R-project.org). Predicted metagenomes were then used as inputs in HUMAnN2 ^24^ to estimate the relative abundances of KEGG Pathways and/or KEGG modules. Based on the KO estimates, relative abundance and coverage of KEGG Pathways was inferred by HUMAnN2. KO information was also used by MinPath ^25^ to infer coverage and relative abundances of KEGG modules, which are manually defined tight, functional units. KEGG Pathways and KEGG modules (KEGG v77.1) data for HUMAnN2 were updated according to publicly available information in IMG ^21^ and KEGG ^22^. Relative abundances and coverages of KEGG modules were represented through circular cladograms generated through GraPhlan.

### Identification of metagenomic biomarkers

We furthered our study through detection of taxonomic clades, KEGG orthologs and metabolic modules that are significantly over/under-represented in the individual oil contaminated environments through statistical analyses carried out on the inferred relative abundances. To this end, the procedure of linear discriminant analysis (LDA) effect size was employed through LEfSe v1.0 ^8^ to identify differentially abundant features that can be used as potential metagenomic biomarkers. For this analysis, the alpha parameter significance threshold for the Krushkal-Wallis (KW) test implemented among classes in LEfSe was set to 0.01 and the logarithmic LDA score cut-off was set to 2.0, due to the relatively small sample size under consideration. All analysis carried out through LEfSe was performed through the Galaxy server ^26^. Additionally, to estimate the associations between taxonomic and functional enrichments in each oil polluted environment, we carried out tests of correlation between abundances for KEGG orthologs and taxonomic clades using a non-parametric test of Spearman’s rank correlation. Detection of significant relationships, defined as a correlation > 0.7 with a p-value < 0.001 and reaching a Benjamini-Hochberg false discovery rate < 0.01 was carried out through the function *corr*.*test* implemented in the R package, *psych* ^27^. Correlations were only computed for oil polluted sites represented by greater than 6 samples. The resultant correlation network was visualized using the interactive platform, Cytoscape v3.4.0 ^28^.

### Detection of microbial interactions

Bacterial interactions in oil contaminated environments was investigated in the present study through non-random bacterial co-occurrence and co-exclusion relationships within individual soil sites. Only polluted sites consisting of more than 4 samples were subjected to deductions of bacterial interactions. mothur implementation of the Sparse Correlations for Compositional data algorithm (SparCC) ^7^, a tool capable of computing significant correlations from compositional data while correcting for the effects of the same, was used to detect significant co-occurrence and co-exclusion patterns. SparCC was run on absolute count OTU tables generated by mothur for each sample, using the command *sparcc* with default settings except a single non-default parameter of permutations=10,000. OTU associations with an absolute SparCC correlation value above 0.6 with p-values < 0.01 were considered statistically significant and incorporated into subsequent network construction. The final network of significant SparCC correlations was built in Cytoscape 3.4.0 ^28^. The nodes in the reconstructed networks represent OTUs participating in robust, statistically significant relationships (both positive and negative), which are in turn portrayed by edges i.e. connections between the nodes.

### Data Availability

16S rRNA amplicon sequencing data generated in this study were deposited in the NCBI Sequence Read Archive (SRA) under accession numbers SRR3168574-SRR3168577. The amplicon sequence data are bundled under NCBI BioProject number PRJNA306989.

## Results

### Bacterial community composition of oil polluted sediments in India

In the present study, we collected oil contaminated samples from sites subjected to regular pollution events in state owned oil refineries at Guwahati and Barhola in the Indian state of Assam to assess the *in situ* bacterial community composition of the same (Table 1). To our knowledge, except for a metagenomic study on oil pipeline microbial populations by Joshi et al. ^29^, this is the first study of its kind to be performed in India. To add diversity to our samples, sampling was carried out from both surface and subsurface soils. 16S rRNA amplicon sequencing data generated by pyrosequencing was further analyzed through mothur, which classified the resulting OTUs into 465 phylotypes. This included 11 phyla, 29 orders and 28 families identified at ≥ 0.5% abundance in at least one of the samples (Fig. 1, Supplementary Table S2). Proteobacteria was the most dominant phylum in all the samples with an average relative abundance of ≥ 50% (Fig. 2). The relative abundance of Proteobacteria (70%) was higher in the Barhola subsurface sample as compared to others (Fig. S1). Acidobacteria was almost as abundant as Proteobacteria in the Noonmati samples (~40%) while lower abundance was observed in the Barhola samples (~10%) (Supplementary Fig. S1). Bacteroidetes and Chlorobi exhibited increase in abundance in the Barhola surface sample when compared to others while Actinobacteria and Chloroflexi were seen to be present in low but consistent abundance across all samples (Supplementary Fig. S1). At the family level, bacterial community composition was much more divergent than at the phylum level with some of them more abundant in particular samples. For instance, Acidobacteraceae (33%), Xanthomonadaceae (10%) and Sphingomonadaceae (7.5%) were detected at higher abundances in Noonmati surface than in others while Koribacteraceae (25%) and Rhodocyclaceaea (~7%) were found to be more enriched in Noonmati subsurface (Fig. 1). Additionally, Hydrogenophilaceae (~9%) and Hyphomicrobiaceae (~4%) exhibited increased abundance in Barhola subsurface compared to other samples (Fig. 1). Chitinophagaceaea, Ectothiorhodospiraceae and Ignavibacteraceae were more enriched in the Barhola samples, while Acetobacteraceae was found in greater numbers in the Noonmati samples (Fig. 1). All samples, except Noonmati surface, showed a high abundance of Comamonadaceae whereas Weeksellaceae and Syntrophaceaea seemed to be specific to Barhola surface (Fig. 1). Families as Sinobacteraceae, Microbacteriaceae and Solibacteraceae were found in fairly consistent abundances across samples (Fig. 1). Overall, at the phylum level the bacterial community composition of Noonmati samples was observed to be more homogenous and less influenced by sampling depth than the Barhola samples (Fig. 1, Fig. 2, Supplementary Fig. S1).

**Figure 1.**
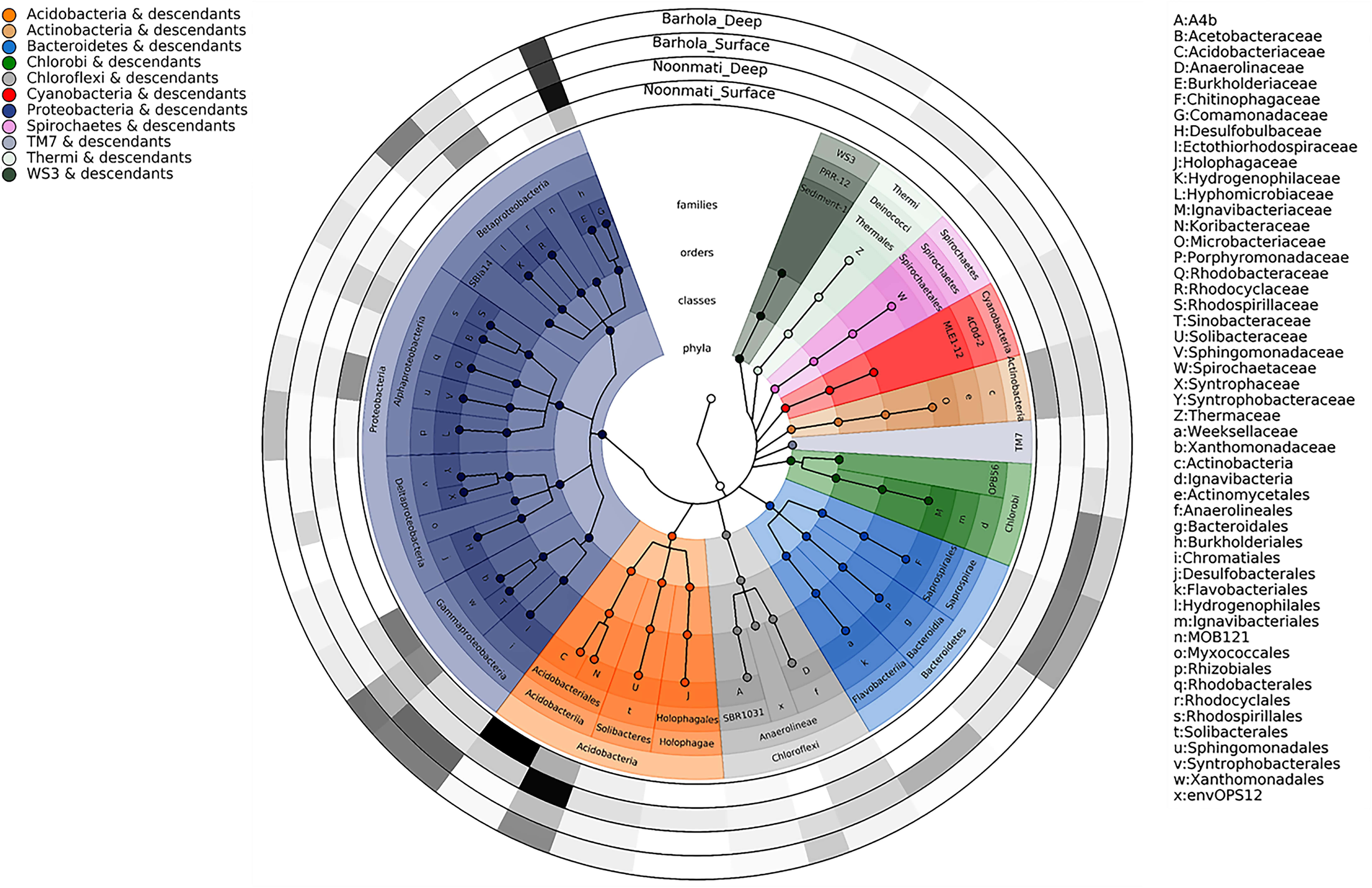
Taxonomic composition of bacterial communities in Noonmati and Barhola oil contaminated soil. Taxonomic cladogram showing all taxa detected at a relative abundance ≥ 0.5% in at least one of the samples. The four rings of the cladogram represent phyla (innermost), class, order and family (outermost) respectively. Circles in the cladogram depict detected taxonomic clades and are colored according to corresponding phyla. Outermost circular rings (external to the cladogram), show rectangular heatmaps depicting abundance of corresponding families in the cladogram. Increasing abundance is represented by increasing opacity.

**Figure 2.**
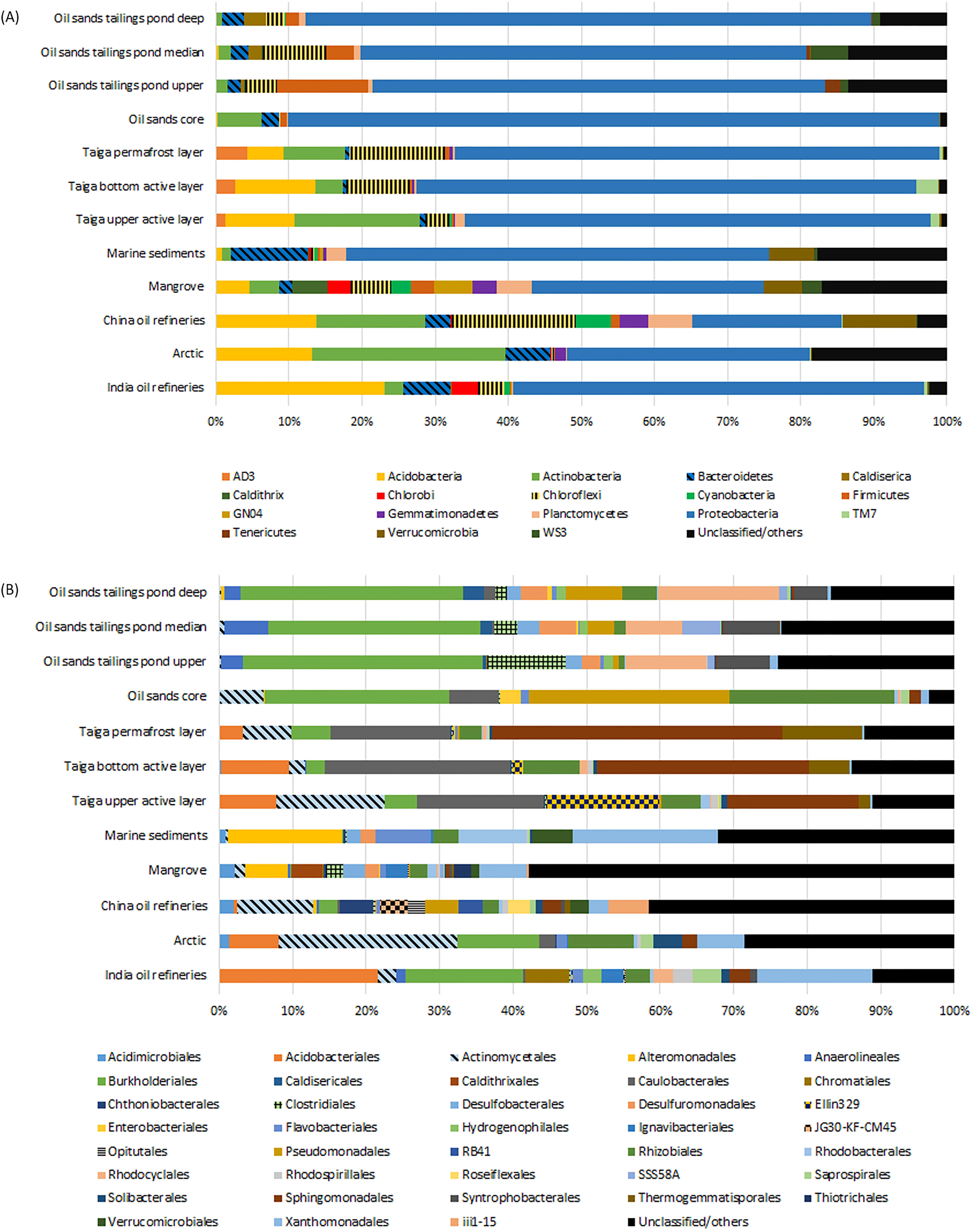
Taxonomic distribution of bacterial communities in oil contaminated environments. Taxonomic clades detected at an average relative abundance ≥ 2% in at least one of 12 oil contaminated habitats, at the phylum level (A), and at the order level (B).

### General characterization of bacterial community composition in petroleum hydrocarbon polluted habitats

Comprehensive characterization of bacterial community composition in hydrocarbon polluted environments was carried out using 61 publicly available and previously validated/published 16S rRNA amplicon sequencing datasets distributed over 11 different habitats (Table 1, Supplementary Table S1) along with 4 datasets generated in this study. mothur analysis of all datasets led to the identification of 18 phyla, 38 orders and 39 families at ≥ 2% abundance in at least one habitat (Fig. 2A, Fig. 2B). Proteobacteria dominated the bacterial community composition at the phylum level with relative abundances ranging from 20-77% across samples (Fig. 2A). Acidobacteria was detected in large numbers in all samples with notably decreased abundances in OSC, OSTPu, OSTPm and OSTPd samples (Fig. 2A, Supplementary Table S1). Actinobacteria and Chloroflexi were consistently identified in all samples with significant increase in A samples, while Bacteroidetes showed higher abundance in DWH and I samples (Fig. 2A, Supplementary Table S1). Similar to our findings, an increase in abundance for the Actinobacteria was reported by Yergeau et al. in diesel contaminated arctic soil biopiles ^30^. Additionally, Chlorobi was detected in high abundance only in M and I samples with increased Gemmatimonadetes abundance identified in A, C and M samples (Fig. 2A, Supplementary Table S1). Verrucomicrobia contribution in microbial community composition was higher in DWH, M and C, while abundance of Firmicutes was higher in OSTP samples and Cyanobacteria in C samples as compared to others (Fig. 2A, Supplementary Table S1). Order level clades with higher abundances detected at ≥ 2% abundance in at least one habitat, tended to be more specific to certain samples. For instance, Acidobacterales had a 21% abundance in I, while Burkholderiales had an average abundance of 30% across OSTP samples and Caulobacerales had an abundance of ~19% in taiga samples (Fig. 2B, Supplementary Table S1). Additionally, Xanthomonadales showed high abundance (15-20%) in I and DWH samples and Actinomycetales dominated A samples with an abundance of 24% (Fig. 2B, Supplementary Table S1). In addition, Alteromonadales (15%) was found in increased abundance in DWH samples, Ellin329 (15%) abundance was highly elevated in Taiga upper active layer (Tu), and Burkholderiales (25%), Pseudomonadales (27%), Rhizobiales (22%) were enriched in OSC (Fig. 2B, Supplementary Table S1). Bacterial families detected at ≥ 2% abundance in a habitat also exhibited preferential sequestration to certain samples. While Caulobacteraceae and Sphingomonadaceae were highly enriched in the taiga samples with an average relative abundance of ~19% and ~29%, Comamonadaceae exhibited a highly elevated mean abundance of 30% in the OSTP samples (Supplementary Table S1). Additionally, Comamonadaceae dominated the I samples bacteriome with an abundance of 15% and contributed 10% of the bacteriome in A samples (Supplementary Table S1). Highly specific increases in relative abundance as compared to other samples included Microbacteriaceae (19%) for A samples, Alteromonadaceae (14%) and Xanthomonadaceae (20%) for DWH samples, and Moraxellaceae (26%) for OSC samples (Supplementary Table S1).

### Similarity in bacterial community structure and detection of taxonomic biomarkers of oil polluted environments

Bray-Curtis similarity scores were inferred from taxonomic data generated by mothur in PAST v3.11 (Table 2) and consequently reduced to a two-dimensional space using NMDS (Fig. 3) for estimation of structural similarity of bacteriomes from petroleum hydrocarbon polluted environments. PERMANOVA tests carried out in PAST showed that taxonomic composition of bacterial communities in the oil polluted environments were significantly varied (*p* = 0.05). However, there were three exceptions. The PERMANOVA results demonstrated that the taiga samples and OSTP samples were not significantly different among themselves (*p* = 0.2-0.9) and that bacteriomes at these sites although separated by depth shared substantial similarity. These observations indicated that unlike large distance spatial separation i.e. geographical isolation, depth or local spatial separation is not a major defining factor for effecting substantial dissimilarity. This is well supported by the Bray-Curtis indices (Table 2) and NMDS plots of the same (Fig. 3) wherein all these samples cluster fairly closely. Additionally, polluted mangrove sediments showed similarity with OSTPm and Tp samples (*p* = 0.057-0.09). Given the very low *p* values these may be aberrations and may have occurred due to preferences, assumptions, and thresholds set in our analysis pipeline. All habitats showed considerable conservation of taxonomic composition within respective samples as described in Table 2. Among these intra-group interactions, OSC samples were indeed clustered in very close proximity (Fig. 3) and exhibited a Bray-Curtis similarity score of 0.85 ± 0.09, which was the highest among all inter and intra-group comparisons (Table 2). Intra-group comparisons of taiga samples showed lowest similarities (Bray-Curtis similarity score 0.45-0.57 ± 0.15) among all habitats, probably due to sampling of source soil from 4 different regions of the China-Russia crude oil pipeline (Table 1, Table 2). Among the inter group comparisons, lowest similarity was observed among M and Tp samples (Bray-Curtis similarity score 0.31 ± 0.02). Apart from the taiga and OSTP samples, which showed inter-group Bray-Curtis similarity score similar to intra-group scores (Table 2), the highest inter-group similarity score of 0.54 ± 0.05 was seen between the relatively related environments of M and DWH.

**Table 2.** Similarities of bacterial community structure within a habitat and between pairs of habitats.

| Habitat | India oil refineries | Arctic | China oil refineries | Mangrove | Marine sediments | Taiga upper active layer | Taiga bottom active layer | Taiga permafrost layer | Oil sands core | Oil sands tailings pond upper | Oil sands tailings pond median | Oil sands tailings pond deep |
| --- | --- | --- | --- | --- | --- | --- | --- | --- | --- | --- | --- | --- |
| India oil refineries | **0.63 ± 0.06** | 0.51 ± 0.07 | 0.39 ± 0.03 | 0.44 ± 0.04 | 0.47 ± 0.04 | 0.41 ± 0.06 | 0.41 ± 0.04 | 0.36 ± 0.05 | 0.48 ± 0.04 | 0.46 ± 0.08 | 0.46 ± 0.08 | 0.48 ± 0.08 |
| Arctic | 0.51 ± 0.07 | **0.72 ± 0.07** | 0.48 ± 0.05 | 0.46 ± 0.03 | 0.42 ± 0.06 | 0.45 ± 0.07 | 0.41 ± 0.05 | 0.39 ± 0.05 | 0.49 ± 0.05 | 0.39 ± 0.05 | 0.39 ± 0.04 | 0.41 ± 0.06 |
| China oil refineries | 0.39 ± 0.03 | 0.48 ± 0.05 | **0.69 ± 0.08** | 0.48 ± 0.02 | 0.40 ± 0.05 | 0.38 ± 0.06 | 0.35 ± 0.05 | 0.34 ± 0.06 | 0.37 ± 0.06 | 0.32 ± 0.02 | 0.35 ± 0.04 | 0.34 ± 0.05 |
| Mangrove | 0.44 ± 0.04 | 0.46 ± 0.03 | 0.48 ± 0.02 | **0.83 ± 0.02** | 0.54 ± 0.05 | 0.34 ± 0.03 | 0.34 ± 0.02 | 0.31 ± 0.02 | 0.36 ± 0.02 | 0.41 ± 0.02 | 0.43 ± 0.04 | 0.41 ± 0.05 |
| Marine sediments | 0.47 ± 0.04 | 0.42 ± 0.06 | 0.40 ± 0.05 | 0.54 ± 0.05 | **0.77 ± 0.09** | 0.35 ± 0.05 | 0.35 ± 0.02 | 0.33 ± 0.05 | 0.43 ± 0.02 | 0.38 ± 0.03 | 0.39 ± 0.03 | 0.42 ± 0.04 |
| Taiga upper active layer | 0.41 ± 0.06 | 0.45 ± 0.07 | 0.38 ± 0.06 | 0.34 ± 0.03 | 0.35 ± 0.05 | **0.52 ± 0.18** | 0.59 ± 0.17 | 0.52 ± 0.18 | 0.45 ± 0.09 | 0.34 ± 0.06 | 0.35 ± 0.05 | 0.37 ± 0.07 |
| Taiga bottom active layer | 0.41 ± 0.04 | 0.41 ± 0.05 | 0.35 ± 0.05 | 0.34 ± 0.02 | 0.35 ± 0.02 | 0.59 ± 0.17 | **0.57 ± 0.12** | 0.55 ± 0.20 | 0.45 ± 0.05 | 0.33 ± 0.04 | 0.34 ± 0.04 | 0.37 ± 0.06 |
| Taiga<br>permafrost<br>layer | 0.36 ±<br>0.05 | 0.39<br>±<br>0.05 | 0.34 ±<br>0.06 | 0.31 ±<br>0.02 | 0.33 ±<br>0.05 | 0.52 ±<br>0.18 | 0.55 ±<br>0.20 | **0.45 ±**<br>**0.22** | 0.42 ±<br>0.10 | 0.32 ±<br>0.06 | 0.33 ±<br>0.05 | 0.36 ±<br>0.08 |
| Oil sands<br>core | 0.48 ±<br>0.04 | 0.49<br>±<br>0.05 | 0.37 ±<br>0.06 | 0.36 ±<br>0.02 | 0.43 ±<br>0.02 | 0.45 ±<br>0.09 | 0.45 ±<br>0.05 | 0.42 ±<br>0.10 | **0.85 ±**<br>**0.09** | 0.45 ±<br>0.05 | 0.45 ±<br>0.04 | 0.56 ±<br>0.10 |
| Oil sands<br>tailings<br>pond upper | 0.46 ±<br>0.08 | 0.39<br>±<br>0.05 | 0.32 ±<br>0.02 | 0.41 ±<br>0.02 | 0.38 ±<br>0.03 | 0.34 ±<br>0.06 | 0.33 ±<br>0.04 | 0.32 ±<br>0.06 | 0.45 ±<br>0.05 | **0.67 ±**<br>**0.12** | 0.64 ±<br>0.09 | 0.63 ±<br>0.13 |
| Oil sands<br>tailings<br>pond<br>median | 0.46 ±<br>0.08 | 0.39<br>±<br>0.04 | 0.35 ±<br>0.04 | 0.43 ±<br>0.04 | 0.39 ±<br>0.03 | 0.35 ±<br>0.05 | 0.34 ±<br>0.04 | 0.33 ±<br>0.05 | 0.45 ±<br>0.04 | 0.64 ±<br>0.09 | **0.56 ±**<br>**0.05** | 0.61 ±<br>0.12 |
| Oil sands<br>tailings<br>pond deep | 0.48 ±<br>0.08 | 0.41<br>±<br>0.06 | 0.34 ±<br>0.05 | 0.41 ±<br>0.05 | 0.42 ±<br>0.04 | 0.37 ±<br>0.07 | 0.37 ±<br>0.06 | 0.36 ±<br>0.08 | 0.56 ±<br>0.10 | 0.63 ±<br>0.13 | 0.61 ±<br>0.12 | **0.61 ±**<br>**0.15** |

**Figure 3.**
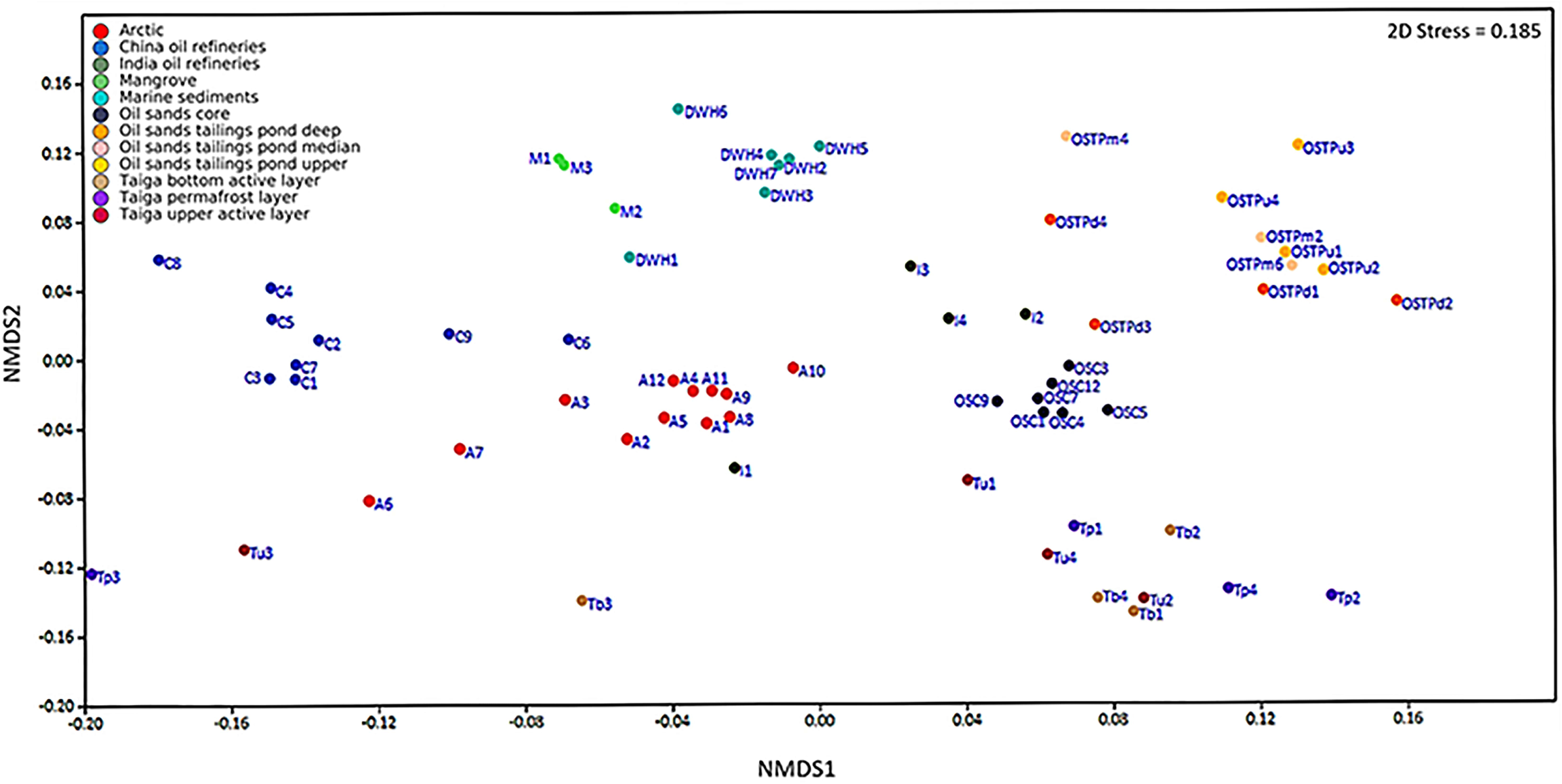
Non-metric multidimensional scaling (NMDS) plot of taxonomic composition of all oil contaminated samples of all habitats. NMDS ordination of 65 oil contaminated samples across 12 habitats was carried out based on Bray-Curtis similarity distances calculated from pairwise taxonomic profile comparisons between all samples. Taxonomic clades present in at least one sample at a relative abundance ≥ 0.5% were used as input. A shorter linear distance between two samples denote greater similarity between the corresponding samples. Samples from 12 environments are depicted by different colors.

To further investigate taxonomic apportionment and detect differentially abundant clades in various oil polluted environments, we compared the abundances of clades detected at an abundance of ≥0.5% in at least 5 samples, at each taxonomic level (Fig. 4). The consequent taxonomic profile inferred for all samples (from domain to species level) was then used by LEfSe to detect metagenomic biomarkers. In all, LEfSe detected 255 differentially abundant taxa including 66 families, 47 genera and 11 species level biomarkers across all habitats (Fig. 4, Supplementary Table S2). The largest number of taxonomic biomarkers were detected for the C samples (68) while the lowest were recorded for both OSTPd and Tu (7). The very low number of detected taxonomic biomarkers for OSTPd and Tu may be a fallout of the comparatively higher bacterial community structure similarity between taiga and OSTP samples than others leading to smaller tally of unique and significantly differential clades. Among the biomarkers detected at the family level, families such as *Acetobacteraceae*, *Rhodospirillaceae*, *Ignavibacteriaceae* and *Chitinophagaceae* were attributed to I samples, *Microbacteriaceae* to A, *Pirellulaceae* and *Planctomycetaceae* to C, *Flavobacteriaceae*, *Rhodobacteraceae,* and *Xanthomonadaceae* to DWH, *Erythrobacteraceae* and *Desulfuromonadaceae* to M, *Rhodocyclaceae* to OSTPd, *Pseudomonadaceae*, *Anaerolinaceae* and *Syntrophaceae* to OSTPm, *Comamonadaceae* and *Geobacteraceae* to OSTPu, *Caulobacteraceae*, *Bradyrhizobiaceae*, and *Hyphomicrobiaceae* to Tb, *Methylobacteriaceae* to OSC, *Thermogemmatisporaceae*, *Alcaligenaceae* and *Sphingomonadaceae* to Tp samples and *Burkholderiaceae*, *Nocardioidaceae*, and *Micrococcaceae* to Tu samples (Fig. 4, Supplementary Table S2). At the genus level, *Phenylobacterium* and *Novosphingobium* were detected as biomarkers for Tp samples, while genera such as *Geobacter*, *Syntrophus*, *Microbacerium*, *Mycobacterium*, *HB2*_*32*_*21*, *Candidatus*_*Koribacter*, *Methylobacterium*, *Caulobacter*, and *Rhodococcus* were attributed as biomarkers for OSTPu, OSTPm, A, C, DWH, I, OSC, Tb, and Tu samples respectively (Fig. 4, Supplementary Table S2). Interestingly, LEfSe detected 19 phylum level biomarkers which indicate that preferential proliferation of bacterial lineages emanating from particular higher level taxa, probably driven by hydrocarbon stress, is possible and may lead to definitive compositional differences between oil polluted habitats. Moreover, candidate phyla such as AC1, WS3 and WS6 were identified as biomarkers for OSTP samples which also underline the uniqueness of these environments (Fig. 4, Supplementary Table S2).

**Figure 4.**
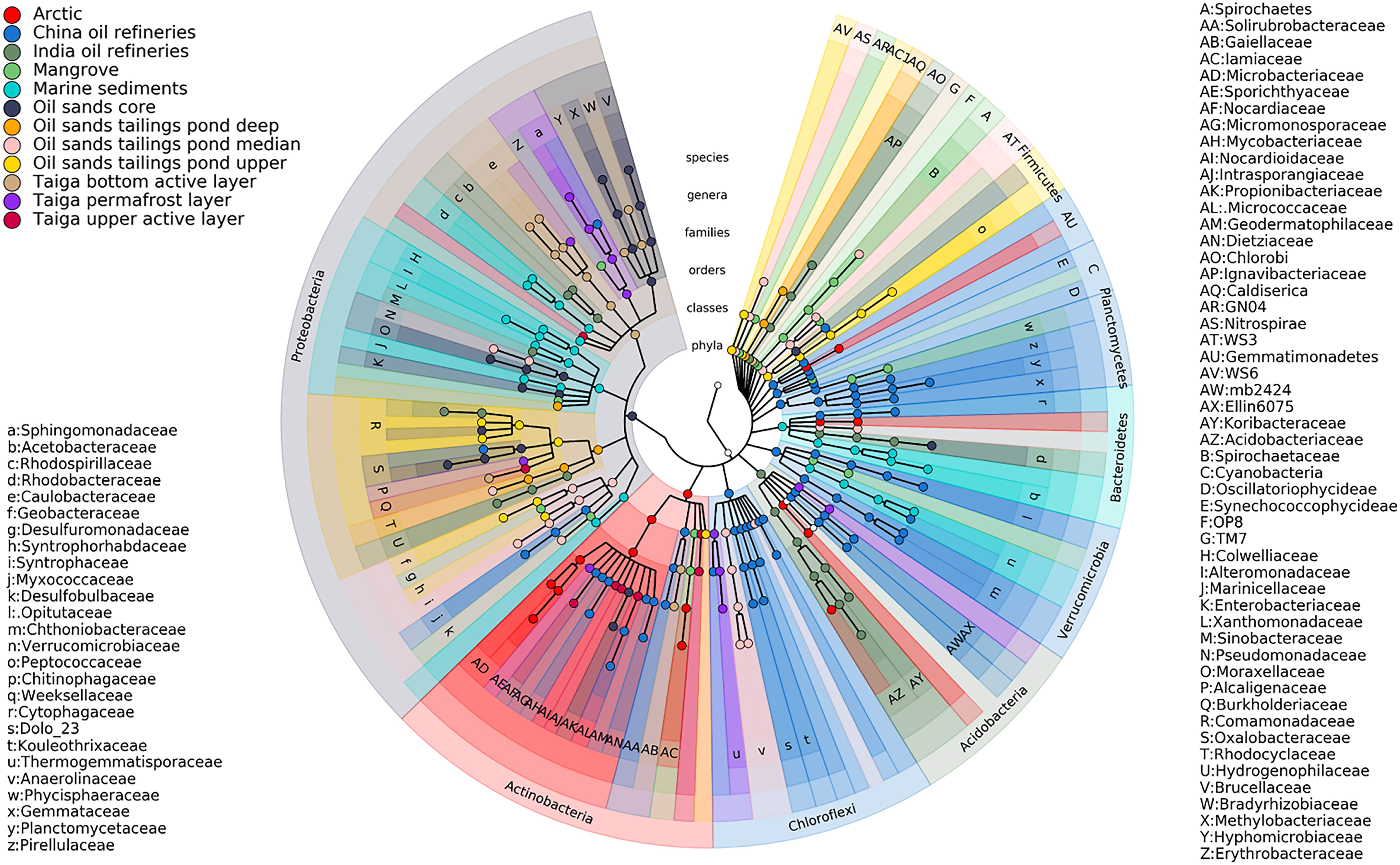
Taxonomic biomarkers of bacterial communities from oil polluted habitats. Cladogram showing all taxonomic clades detected at a relative abundance ≥ 0.5% in at least five samples across all habitats. These were used as inputs for LEfSe. Seven rings of the cladogram represent phylum (innermost), class, order, family, genus and species (outermost), respectively. Enlarged circles represent differentially abundant taxa detected as taxonomic biomarkers and are colored corresponding to the individual soil habitat wherein they are over-represented among 12 oil polluted ecosystems (see legend).

### Metabolic characterization and functional biomarkers of oil contaminated environments

For understanding the metabolic potential of oil polluted environments and identifying differentially abundant functional features, metagenomes were predicted by PICRUSt using the 16S rRNA gene amplicon data generated. Predicted proteins were classified as KEGG orthologs (KOs) resulting in the identification of 7020 KOs across all samples. Metabolic reconstruction of metagenomes predicted by PICRUSt was carried out in HUMAnN2, which detected 585 KEGG modules across all samples. Among these functional modules, 19 functional modules were present across all samples at a coverage of >90% and were identified as core modules (Fig. 5, Supplementary Fig. S2, Table 3, Supplementary Table S4). Most of the core modules identified are essential for sustenance of prokaryotic life in the environments, such as translation (M00178), central carbon metabolism (M00149), ATP synthesis (M00153, M00157) and nucleotide and amino acid metabolism (M00005, M00020). Rest of the core modules identified were found to be involved in various kinds of transport systems for cations, nutrients and peptides including iron, phosphate, nickel, and amino acids (M00188, M00222, M00223, M00236, M00237,M00239, M00240, M00250, M00254, M00255, M00256, M00258, M00320) (Fig. 5, Supplementary Fig. S2, Table 3, Supplementary Table S4). This is important since these resources are generally present in limiting quantities in nature and often determine the survival and proliferation of microbes in the environment. Additionally, transport systems for lipopolysaccharide (LPS), a principal component of the gram-negative bacterial cell wall, were also understandably identified as core modules and included KEGG functional modules for export of LPS across both cytoplasmic (M00250) and outer membranes (M00320) (Fig. 5, Supplementary Fig. S2, Table 3, Supplementary Table S4). Furthermore, 56 differently covered functional modules were detected across all oil contaminated samples (Fig. 5, Supplementary Fig. S2, Supplementary Table S4). Among these, five modules were completely covered in only one sample while being absent in all others (Fig. 5, Supplementary Fig. S2, Supplementary Table S4). This included structural complexes for Manganese/Iron transport (M00243), bacterial proteasomes (M00342) and putative aldouronate transport (M00603), all of which were completely covered only in the C samples (Fig. 5, Supplementary Fig. S2, Supplementary Table S4). This indicates that bacteria in the C site are better equipped for transport of metallic cations, peptide utilization and uptake of plant derived aldouronates than other sites. Furthermore, the presence of a complete complement of D-Xylose transport system (M00215) in the C site also reaffirms bacterial access to hemicellulosic plant material at this site (Fig. 5, Supplementary Fig. S2, Supplementary Table S4). Additionally, glutamate transport system (M00233) was completely covered at only the A site, and RstB-RstA stress response two component system (M00446) at the OSC site (Fig. 5, Supplementary Fig. S2, Supplementary Table S4). The bacteria at A site, thus are extremely capable of utilizing glutamate for growth, while resident bacteria at OSC are better furnished with stress response mechanisms critical in environmental adaptation and survival.

**Figure 5.**
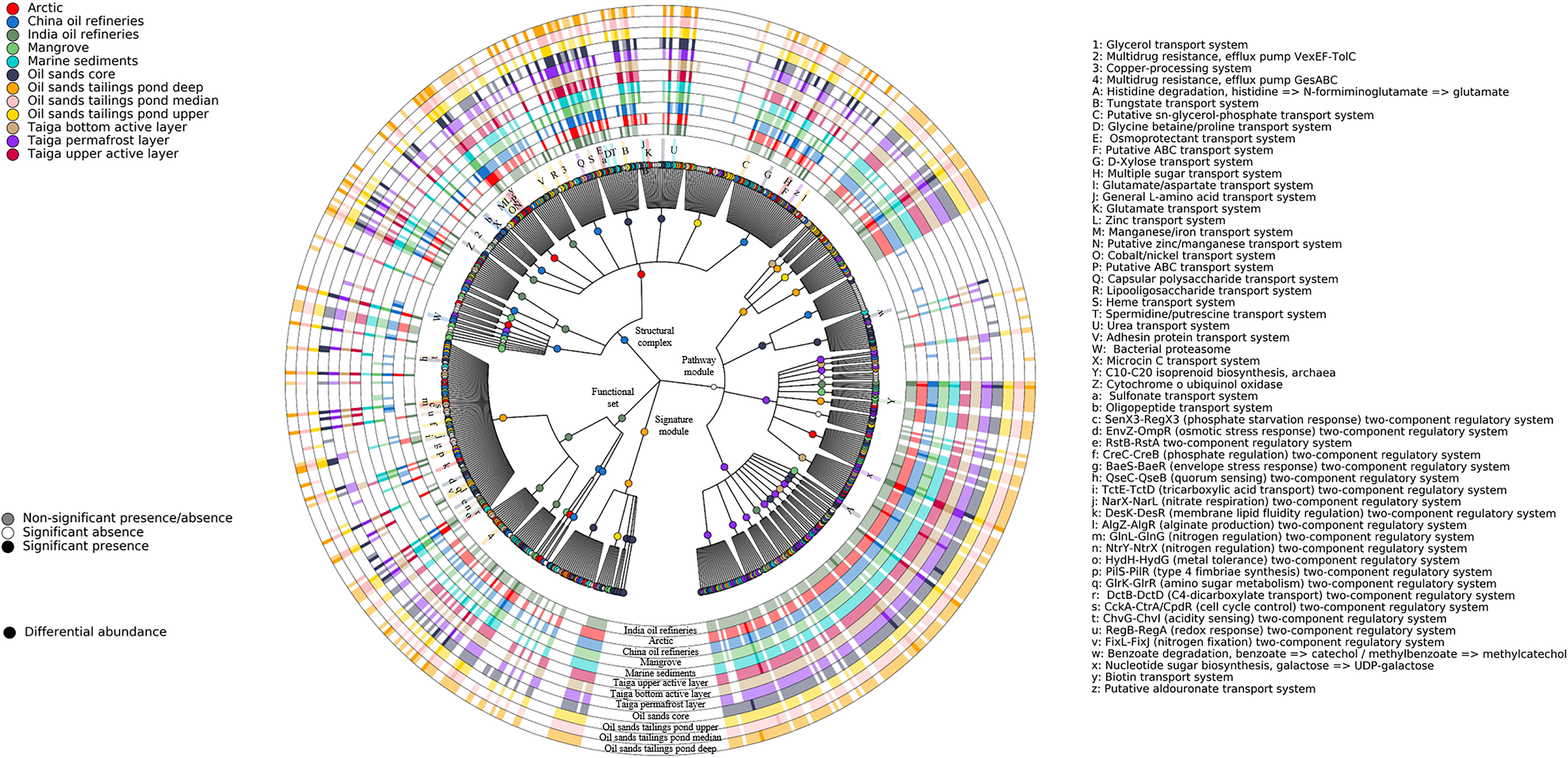
Metabolic reconstruction of metagenomes from oil polluted habitats. Cladogram showing KEGG BRITE hierarchical structures denoted by innermost four rings as inferred against detected KEGG metabolic modules for all oil contaminated samples. Outermost ring represents KEGG functional modules that have been detected in at least one of the 65 PICRUSt predicted metagenomes as reconstructed by HUMAnN2. Over-represented metabolic modules inferred by LEfSe are depicted by enlarged circles and are colored corresponding to the oil contaminated habitat they have been identified to be differentially abundant in. Outermost rings (external to the cladogram) depict heat-maps representing significantly or non-significantly present/absent modules across all oil polluted environments. Presence is represented by ≥ 90% coverage and absence by ≤ 10% coverage of KEGG module as estimated by HUMAnN2. Varied modules are labeled and included in legend.

**Table 3:** Core modules shared between habitats as detected by HUMAnN2.

| **Module ID** | **Definition of modules in KEGG** |
| --- | --- |
| M00005 | PRPP biosynthesis, ribose 5P => PRPP |
| M00020 | Serine biosynthesis, glycerate-3P => serine |
| M00149 | Succinate dehydrogenase, prokaryotes |
| M00153 | Cytochrome d ubiquinol oxidase |
| M00157 | F-type ATPase, prokaryotes and chloroplasts |
| M00178 | Ribosome, bacteria |
| M00188 | NitT/TauT family transport system |
| M00222 | Phosphate transport system |
| M00223 | Phosphonate transport system |
| M00236 | Putative polar amino acid transport system |
| M00237 | Branched-chain amino acid transport system |
| M00239 | Peptides/nickel transport system |
| M00240 | Iron complex transport system |
| M00250 | Lipopolysaccharide transport system |
| M00254 | ABC-2 type transport system |
| M00255 | Lipoprotein-releasing system |
| M00256 | Cell division transport system |
| M00258 | Putative ABC transport system |
| M00320 | Lipopolysaccharide export system |

In addition to differently covered functional modules, 414 KEGG modules were detected to be differentially abundant in at least one of the 12 contaminated environments (Fig. 5, Supplementary Fig. S2, Supplementary Table S3). The largest number of differentially abundant modules were attributed to the OSC samples (70) while the least (8) were attributed to the OSTPm samples (Fig. 5, Supplementary Fig. S2, Supplementary Table S3). The detection of a higher number of differentially abundant modules in the OSC samples is possibly due to its highly extreme environment as compared to other samples, leading to sequestration of a large number of convenient functions to optimize the use of available resources and counteract distinct environmental stress conditions. On the contrary, similar to the result for taxonomic biomarkers, the least number of differential functional modules were detected in an OSTP sample (OSTPm), with the penultimate spot being taken by Tu (13). As explained above, this is not surprising since both taiga and OSTP samples share comparatively greater similarity between their habitats leading to an overlap of functional capabilities and hence, fewer unique and over-represented functional modules (Fig. 5, Supplementary Fig. S2, Supplementary Table S3). Most of the modules for metabolism of aromatic hydrocarbons such as xylene degradation (M00537), toluene degradation (M0053), benzoate degradation (M00540 and M00551), salicylate degradation (M00638) and catechol ortho-cleavage (M00568) were also significantly associated with the OSC samples (Fig. 5, Supplementary Fig. S2, Supplementary Table S3). A number of structural complexes implicated in photosynthesis were found to be differentially abundant in C samples which included Photosystems I and II (M00163, M00161), the cytochrome b6f complex (M00162) and NADP(H): Quinone oxidoreductase for chloroplasts and cyanobacteria (M00145) (Fig. 5, Supplementary Fig. S2, Supplementary Table S3). Additionally, a plethora of amino acid biosynthesis modules were detected as functional biomarkers in the taiga samples. For example, three different modules for lysine biosynthesis (M00525-M00527), and one each for threonine, methionine and cysteine biosynthesis (M00018, M00017, M00021) were significantly abundant in Tb samples while modules for valine/isoleucine, phenylalanine, tyrosine, leucine and isoleucine biosynthesis (M00019, M00024, M00026, M00040, M00432, M00535, M00570) were over-represented in Tp samples (Fig. 5, Supplementary Fig. S2, Supplementary Table S3). The taiga samples also exhibited an over-representation for modules involved in the biosynthesis of vitamins and cofactors as heme, pantothenate, ubiquinone, tetrahydrofolate, thiamine and ascorbate (M00127, M00129, M00121, M00119, M00128, M00126) (Fig. 5, Supplementary Fig. S2, Supplementary Table S3).

Overall, all the sites were found to harbor a variety of differentially abundant modules dedicated to the transport of saccharide, polyols, peptides, metallic cations, vitamin, amino acid, mineral ions, organic ions, lipid and phosphate underlining the large genetic investment of resident bacteria in the processing of environmental information specific to the said site (Fig. 5, Supplementary Fig. S2, Supplementary Table S3). However, while differentially over-represented transport systems for saccharides, polyols and lipids were almost ubiquitously detected, significantly associated transport systems for other substrates as phosphates, amino acids, peptides and organic ions were restrained to certain sites. This may indicate differential availability of these nutrients resulting in preferential dependence on certain substrates acquired from the environment and also reaffirms the characteristically different nature of the environments under consideration. A large number of differentially abundant biosynthetic pathways for sugars, amino acids and vitamins were also detected along with a great diversity of two component systems catering to a range of functions such as stress and redox response, quorum sensing, chemotaxis and heavy metal tolerance across all sites (Fig. 5, Supplementary Fig. S2, Supplementary Table S3). Additionally, some modules for atypical energy metabolism as denitrification, dissimilatory nitrate reduction and dissimilatory sulfate reduction were also detected to be differentially abundant and may be important biomarkers for the corresponding sites due to their contribution in bacterial respiration (Fig. 5, Supplementary Fig. S2, Supplementary Table S3). Finally, several modules describing microbial resistance to antibiotics and antimicrobial peptides were detected to be over-represented at all sites (Fig. 5, Supplementary Fig. S2, Supplementary Table S3). This is probably due to the method of ancestral state reconstruction used by PICRUSt for genome prediction, that leads to these genes being predicted for consequent metagenomes if input 16S rRNA data includes hits from bacteria known to have antimicrobial resistance genes. The possession and even expression of these genes probably will not have a significant selective advantage in environments already undergoing natural selection due to oil pollution.

### Associations between bacterial taxa and metagenomic gene families

Correlations between bacterial abundance and functions enriched at different sites were evaluated following a statistical strategy similar to the approach described by Segata et al. ^31^. The results indicated strong and significant associations between a number taxonomic clades and gene families predicted by PICRUSt (Supplementary Fig. S3). A subset of these significant correlations included strong associations between previously detected taxonomic biomarkers and over-represented KOs for each site, which further confirmed the identified taxonomic biomarkers. For example, photosynthetic structural complex genes *cpeA* (K05376) and *psb28-2* (K08904), found to be differentially abundant in C samples exhibited strong positive association with an over-represented cyanobacterial order, Oscillatoriophycideae (Spearman correlation > 0.7, P-value < 0.001) (Supplementary Fig. S3). Additionally, an array of genes related to polycyclic aromatic hydrocarbon degradation such as *nidABD*, *phdFGIEK*, and *phtAaBC* (K11943-48, K18251, K18255-57, K18275) were differentially abundant in A samples and also significantly positively correlated to known polyaromatic hydrocarbon degrader and taxonomic biomarker *Mycobacterium* (Spearman correlation > 0.7, P-value < 0.001) ^32^ (Supplementary Fig. S3). In other observations, *Methylobacterium* showed positive correlation with a number of genes associated with the transport of sugars, saccharides and amino acids such as *ggtB-D* (K10232-34), *cebE-G* (K10240-42), *chvE* (K10546), *gguA-B* (K10547-48), and *gluA-D* (K10005-08) in arctic samples (Spearman correlation > 0.7, P-value < 0.001) (Supplementary Fig. S3). Hydrocarbon degradation genes like *pcaG* (K00448), *bbsH* (K07546) and *pcaL* (K14727) were significantly correlated to class Actinobacteria in a positive manner in the same samples (Spearman correlation > 0.7, P-value < 0.001) (Supplementary Fig. S3). In the DWH sample, Colwelliaceae exhibited positive correlations with both anaerobic C4-dicarboxylate transporter (*dcuB;* K07792) and 2-oxopent-4-enoate/cis-2-oxohex-4-enoate hydratase (*bphH*, *xylJ*, *tesE;* K18820), an enzyme implicated in oligosaccharide metabolism (Spearman correlation > 0.7, P-value < 0.001) (Supplementary Fig. S3). Additionally, genus *HB2.32.21*, associated positively with a number of genes involved in alginate production (*alg44*, *algJXKFE;* K19291-3, K19295-6, K16081), flagellar synthesis/chemotaxis (*qseC;* K07645) and aminobenzoate metabolism gene regulation (*feaR*; K14063) (Spearman correlation > 0.7, P-value < 0.001) (Supplementary Fig. S3). Acidobacteria however, was found to be negatively correlated with the *alkB1-2* gene (K00496) coding for alkane-1-monooxygenase (Spearman correlation < - 0.7, P-value < 0.001) (Supplementary Fig. S3). In OSC samples, positive correlations were detected between *Methylobacterium* and genes involved in furfural degradation (*hmfABCDEF;* K16874-80) and benzoate degradation (*aliAB*, *badL*; K04116-17, K07536) (Spearman correlation > 0.7, P-value < 0.001) (Supplementary Fig. S3). *Methylobacterium*, although an aerobe ^33^, has been shown to possess anaerobic benzene degradation genes in the genome annotation for *Methylobacterium extorquens* PA1 in KEGG (http://www.genome.jp/kegg-bin/show_pathway?mex01220). Furthermore, a number of two-component systems (TCS) showed strong positive association with *Acinetobacter* and Enterobacteriaceae. *Acinetobacter* was positively correlated with the enrichment of RstA/RstB stress response TCS (K07639, K07661), while Enterobacteriaceae showed affirmative relationships with the aerobic stress response sensor kinase ArcB (K07648) and nitrate/nitrite response regulator NarP (K07685) (Supplementary Fig. S3).

To further understand the association of bacterial clades with gene families specifically with respect to hydrocarbon degradation, we categorized all taxa contributing to the abundance of genes known to be involved in hydrocarbon degradation at the family and genus level (Supplementary Fig. S4). The results showed that differences existed between major contributors to the abundance of particular hydrocarbonoclastic genes at different sites. For example, abundance for alkane-1-monooxygenase (K00496) was contributed mainly by Alteromonadaceae in DWH samples, Comammonadaceae in I, Mycobacteriaceae and Nocardiaceae in C, Propionibacteriaceae in OSC, and a mixture of Acetobacteraceae, Mycobacteriaceae, Nocardiaceae and Rhodospirillaceae in the taiga samples (Supplementary Fig. S4). Similarly, for protocatechuate-4,5-dioxygenase (K04100-01), Alteromonadaceae were again the major contributors for DWH samples, Comamonadaceae and Methylobacteriaceae for OSC, Rhodocyclaceae for I, Rhodocyclaceae and Comamonadaceae in OSTP, and Comamonadaceae and Bradyrhizobiaceae for taiga samples (Supplementary Fig. S4). These differences in patterns observed at the family level, were even more stark at higher resolutions i.e. genus level, thus effectively differentiating such metagenomic contributors from site to site. This was best demonstrated for the hydrocarbonoclastic gene catechol-1,2-dioxygenase (K03381), for which Alteromonadaceae was found to be the most dominant contributor in both DWH and M samples (Fig. S4). However, at the genus level, it was seen that while *HB2-32-21* was the dominant effector organism in DWH samples, *Marinobacter* was the largest metagenomic contributor for K03381 in the M samples (Supplementary Fig. S4).

### Bacterial interactions in oil polluted environments

To further understand complex ecological relationships in oil polluted environments, bacterial association networks were deduced from estimated taxonomic profiles. For our study, we concentrated on individual oil polluted habitats, i.e. Arctic, China oil refineries, oil sands core and so on. The resulting bacterial correlation networks, inferred at or above the species level, constituted 186 significant relationships among 115 phylotypes (P < 0.01) (Fig. 6). Among the associations deduced to be significant, 72.58% were detected to share positive correlations while the rest shared antagonistic relationships. Almost half of the co-occurrence patterns identified (46%) were observed between bacteria of the same phyla while more than three-quarters of all negative correlations (78%) were detected between bacteria belonging to distinct phyla (Fig. 6). Thus, our results from the inferred bacterial correlation networks indicated that, co-occurrence of phylotypes was closely related to sharing of evolutionary lineage. For example, in the OSC habitat, phylotypes belonging to proteobacterial family Oxalobacteraceae shared positive pairwise correlations with Moraxellaceae and Enterobacteriaceae phylotypes, both of which belong to phylum Proteobacteria (Fig. 6). Additionally, similar co-occurrence patterns were observed between phylotypes attributed to families belonging to the order Actinomycetales in the C samples. Positive pairwise associations were observed in C samples between phylotypes from families Micrococcaceae and Nocardioidaceae, Intrasporangiaceae and Mycobacteriaceae with Solirubrobacteraceae, and Gaiellaceae and Geodermatophilaceae with Microbacteriaceae, all of which belong to order Actinomycetales (Fig. 6). Furthermore, genera *Arthrospira* and *Phenylobacterium,* both of which belong to family Caulobacteraceae, co-occurred in the Tu samples (Fig. 6). Conversely, bacteria without evolutionary commonalities tended to be negatively correlated. For example, in DWH samples, antagonistic relationships were observed between phylotypes belonging to family Flavobacteriaceae from phylum Bacteroidetes and proteobacterial families Desulfuromonadaceae and Desulfobulbaceae (Fig. 6). Similarly, mutual exclusion was observed between phylotypes belonging to family Weeksellaceae of phylum Bacteroidetes and Xanthomonadaceae of phylum Proteobacteria in I samples. Additionally, negatively correlated associations were observed between phylotypes belonging to genera Pelotomaculum and Thiobacillus in OSTPu samples, the former of which belongs to phylum Firmicutes and the latter to phylum Proteobacteria (Fig. 6).

**Figure 6.**
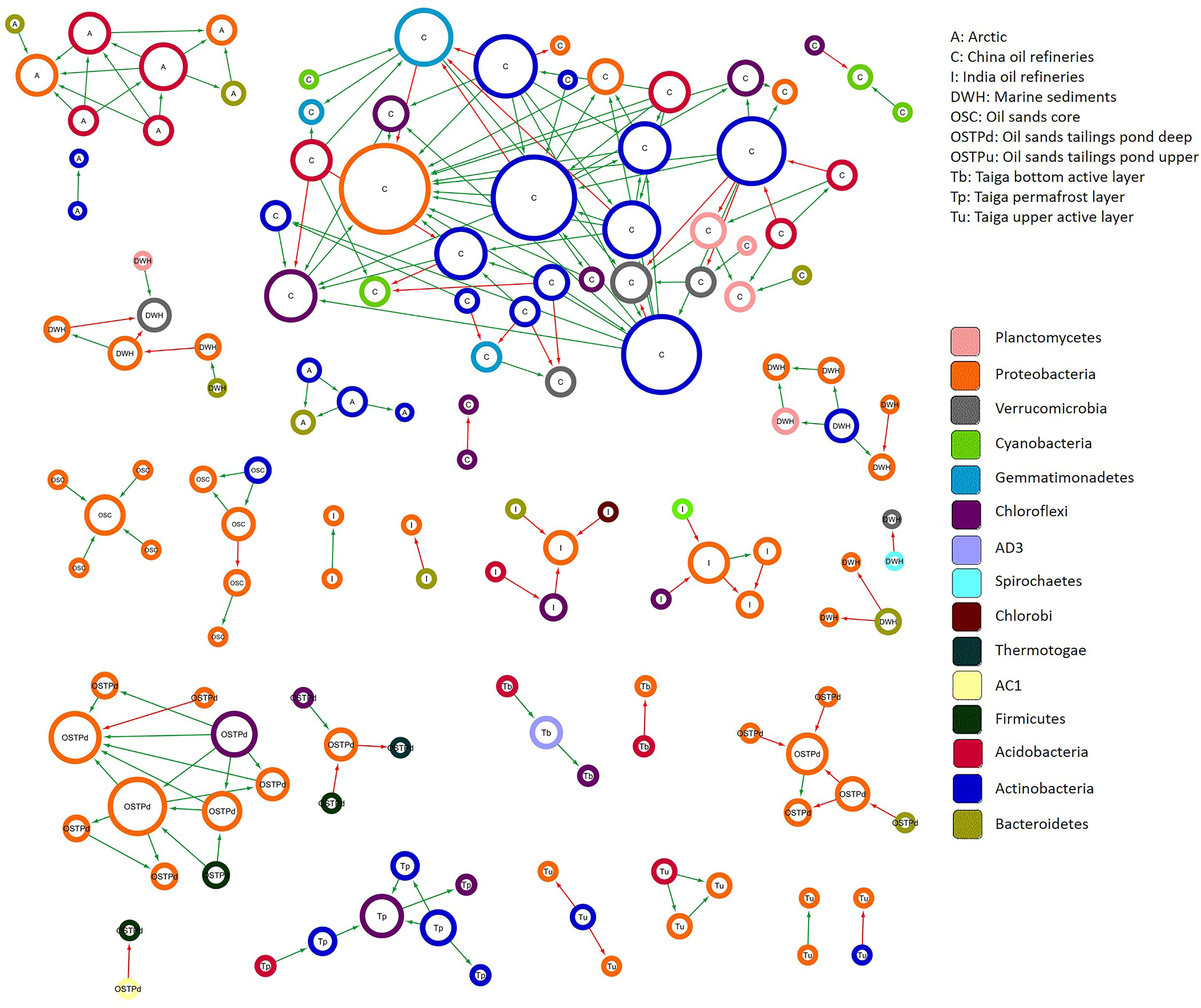
SparCC network plot of global microbial interactions in individual oil polluted habitats. Significant bacterial associations captured by SparCC (*p*-value < 0.01) with an absolute correlation magnitude of ≥ 0.6 are presented. Nodes represent detected phylotypes (OTU clustered at 97% similarity) involved in either significant co-occurrence (green edges) or co-exclusion (red edges) relationships. Border coloration depicts taxonomic affiliation of nodes at the phylum level. Node size is proportional to the connectivity of the node (both positive and negative relationships).

## Discussion

The advent of next-generation sequencing (NGS) technologies have revolutionized investigative approaches into microbial processes. This has led to re-exploration of well-known microbial processes as the nitrogen cycle ^34^, methane metabolism ^35^, sulfur cycle ^36^, heavy metal remediation and petroleum bioremediation ^37^ along with examination of exotic and extreme environments such as deep-sea hydrothermal vents ^38^, cold deserts as Antarctica ^39^ and remote cave systems ^40^. During this time, a large amount of work has been done on the microbiology of hydrocarbon degradation using NGS technologies as well ^41^. Most of these studies employed 16S rRNA based amplicon sequencing while some used metagenomic shotgun sequencing for their enquiries. Although some of these studies have concentrated on prediction of potential biomarkers for oil pollution in certain environments ^42,43^, no investigative effort has been undertaken to use the large amounts of data generated in oil pollution studies across the world to review, validate and further these studies. In the present study, we report bacterial diversity in oil contaminated soil collected from north-eastern India and describe taxonomic and functional characteristics of oil polluted environments across the world in order to understand the differences and similarities that exist between them. Additionally, we infer a large number of potential biomarkers, both taxonomic and functional, along with co-occurrence networks, which provide new insights into the process of oil bioremediation including taxa and metabolic pathways critical to survival in different oil polluted ecosystems. To this end, we have used 65 16S rRNA datasets from different studies across the world (Table 1, Supplementary Table S1), including 4 datasets generated in this study, and carried out robust *in-silico* analysis with recently developed bioinformatics tools to compare and contrast the same. The principal features and findings of our study are discussed below.

### Taxonomic and functional features of oil contaminated soils in India

To unravel the bacterial community structure, functional characteristics and complex inter-relationships in oil contaminated environments in India, crude oil polluted samples were collected from two oil refineries in north-east India. 16S rRNA amplicon sequencing analysis of the said samples revealed an overall predominance of the metabolically versatile phylum Proteobacteria (Fig. 2, Supplementary Fig. S1), as has been reported previously in other oil polluted sites ^44^. Additionally, phylum Acidobacteria was also detected to be prevalent in these samples and was identified as a biomarker as well (Fig. 2, Supplementary Fig. S1, Fig. 4, Supplementary Table S2). Further identification the acidobacterial genera *Edaphobacter* and *Candidatus_Koribacer* as biomarkers underlines the importance of the acidobacterial lineage in the oil contaminated India soils and also indicates a slightly acidic environment (Fig. 4, Supplementary Table S2). *Methylibium petroleiphilum* was also detected as a biomarker and contributed significantly to the hydrocarbon degradation capabilities at these sites (Fig. 4, Supplementary Table S2, Supplementary Fig. S4). *Methylibium petroleiphilum,* an aerobic bacterium, has previously been reported to degrade hydrocarbons as methyl tert-butyl ether ^45^. Moreover, anaerobic genera as the sulfate dissimilating *Thiobacillus* and the photoautrophic *Ignavibacteriaceae*, were also identified as biomarkers (Fig. 4, Supplementary Table S2). These observations, along with the detection of differentially abundant KEGG modules as dissimilatory sulfate reduction, sulfate => H2S (M00596) and NarX-NarL (nitrate respiration) two-component regulatory system (M00471; also found to be present with complete coverage) (Supplementary Fig. S2, Supplementary Table S3, Supplementary Table S4) indicate that anaerobic processes and taxa play a major role in the oil contaminated India soils. Simultaneously, the identification of over-represented aerobic pathways such as formaldehyde assimilation, serine pathway (M00346) (Supplementary Fig. S2, Supplementary Table S3) and differentially abundant aerobic taxa as *Methylibium* and Chitinophagaceae (Supplementary Fig. 4, Supplementary Table S2) however indicate that these environments may be predominantly aerobic but partially anaerobic or microaerophilic. Over-representation of Chitinophagaceae in these environments also indicate possible availability of chitin as a carbon source ^46^. NMDS ordination of taxonomic profiles and estimation of Bray-Curtis similarity indices for I samples show that they share much more similarity amongst themselves than with other sites (Fig. 3, Table 2). I samples showed highest similarity with A samples, which may be due to the enrichment of Acetobacteraceae and Comamonadaceae in both the sites possibly due to oil contamination (Fig. 3, Table 2). This is well demonstrated in metagenomic contributions of both families to hydrocarbonoclastic genes (Supplementary Fig. S4).

However, whereas both the families are the consistently major contributors for hydrocarbon degrading genes across all I samples, other families are shown to also contribute hydrocarbonoclastic capabilities to samples in A sites (Supplementary Fig. S4). Furthermore, samples collected from India oil refineries show extensive adaptation to oil pollution as revealed by the detection of KOs for known hydrocarbonoclastic genes such as alkane-1-monooxygenase, protocatechuate-3,4-dioxygenase, catechol-1,2-dioxygenase and protocatechuate-4,5-dioxygenase (Supplementary Table S3, Supplementary Fig. S4) along with over-represented hydrocarbon degradation genes such as homogentisate-1,2-dioxygenase and the toluene monooxygenase system (*tmoABCDEF* / *tbuA1A2BCUV*/*touABCDEF*) (data not shown). Bacterial interaction networks inferred by SparCC show a high proportion of co-exclusion relationships, most of which exist between taxa belonging to different evolutionary lineages (Fig. 6). Competitive interactions were observed between aerobic and anaerobic taxa, which indicates co-existence in relative proximity with competition for resources and furthers our inference of a possibly microaerophilic or partially anaerobic oil polluted environment (Fig. 6). One of only two positive correlations was found to be shared between *Methylibium* and *Parvibaculum*, which may endow *Methylibium* with a competitive edge and facilitate its enrichment in the I samples (Fig. 6).

### Validation of bioinformatic pipeline

To our knowledge this is the only study that has congregated existing 16S rRNA NGS data generated during experiments on hydrocarbon pollution in different habitats around the world to deduce possible biomarkers and associated bacterial characteristics and interactions. The bioinformatics pipeline we designed to analyze this data employed PICRUSt, which is a recently developed tool that uses 16S rRNA data to predict metagenomes for corresponding samples along with LEfSe which predicts potential biomarkers and HUMAnN2 for metabolic reconstruction of PICRUSt predicted metagenomes. It is to be noted however, that KEGG orthologs and KEGG module databases for PICRUSt and HUMAnN2 were meticulously updated (previously PICRUSt KEGG databases included KOs only up to K15039 and HUMAnN had a KEGG module database represented only up to M00378) to include currently available definitions of KEGG functional modules and represent the metabolic terrain of petroleum hydrocarbon contaminated habitats in totality, especially with respect to hydrocarbon degradation, functional modules for which were absent in the original databases. To confidently interpret and infer our results, we validated our findings in both taxonomic and functional aspects. For example, a complete convergence of conclusion was observed when comparing our inferred taxonomic compositions and biomarkers with the findings of Mason et al. ^47^ for the marine sediments samples. Our analysis of the marine sediment samples identified a highly dominant Gammaproteobacterial genus, HB2-32-21 (Greengenes OTU ID 248394) belonging to the family Alteromonadaceae (Table S2), which was detected as a taxonomic biomarker (Fig. 4) and also contributed significantly to the abundance of hydrocarbon degradation genes at the site (Supplementary Fig. S4). Additionally, Colwelliaceae and Rhodobacteraceae were also detected as over-represented taxonomic biomarkers at the Macondo oil contaminated DWH sample sites (Fig. 4, Supplementary Table S2) with the latter contributing largely abundance of the alkane degrading enzyme, alkane-1-monooxygenase (Supplementary Fig. S4). All these observations, are extremely consistent with the findings of Mason et al. ^47^ in the original article and furthers their study providing new insights. Moreover, we found important similarities between conclusions inferred by An et al. ^17^ and our study, regarding the oil sands core datasets. In the original study by An et al. ^17^, the oil sands core was deduced as an aerobic environment with limited oxygen ingress in specific regions leading to regional anaerobiasis. This theory of intermittent oxygen infusion in sections of the oil sands core was strongly supported by the detection of both aerobic and anaerobic pathways of hydrocarbon degradation in the oil sands core. For example, in the oil sands core samples we detected differentially abundant KEGG modules for aerobic degradation of different hydrocarbons as xylene, benzoate, toluene and cumate including metabolism of corresponding intermediates as salicylate and catechol (M0537-40, M00568, M00638) (Supplementary Fig. S2, Supplementary Table S3) ^48^ alongside a module implicated in anaerobic degradation of benzoate (M00551) (Supplementary Fig. S2, Supplementary Table S3) ^49^. The larger number of aerobic hydrocarbonoclastic modules compared to anaerobic modules therefore further validated our bioinformatic pipeline for consequent interpretation of findings in the present study.

### Metabolic reconstruction of oil polluted metagenomes reveals important functional pathways in petroleum hydrocarbon contaminated habitats

In order to understand the functional landscape of each oil polluted environment, metagenomes were predicted by PICRUSt from 16S rRNA data and metabolic modules detected using HUMAnN2. We identified 19 core modules which were present across all habitats with a coverage of > 90%. Most of these are involved in processes central to survival of bacteria in the environment. Furthermore, in order to identify preferential genetic investments among resident bacteria at each habitat differentially abundant KOs and KEGG modules were detected through LEfSe. Consequently, we analyzed over-represented KOs and KEGG modules across all habitats to identify broad metabolic signatures that may be indicative of important areas of genetic expenditure, especially outside hydrocarbon degradation. As a result, we identified a number of differential functional pathways dedicated to transport of certain sugars or lipids, biosynthesis of particular biomolecules, stress response, quorum sensing, metabolism of polysaccharides, assimilation and respiration of sulphur and/or nitrogen compounds besides hydrocarbon degradation, across all sites. For example, a large number of putrescine transport complexes (M00193, M00299, M00300) a transport system for arginine/ornithine (M00235) were detected to be differentially abundant for the DWH samples (Fig. 5, Supplementary Fig. S2, Supplementary Table S3). This sequestration of putrescine transporters along with ornithine, which is readily converted by ornithine decarboxylase to putrescine indicates a significant dependence of marine bacteria at an oil polluted site on putrescine. This can be explained by the crucial role putrescine plays in bacteria as an osmoprotectant ^50^, and therefore its prevalence in a marine oil polluted environment. Similarly, availability and possible use of carbon sources besides hydrocarbons was apparent in the C samples. The differential presence of a complete complement of D-xylose transport system (M00215) and a putative aldouronate transport system (M00603) along with the over-representation of KEGG module M00014 (Glucuronate pathway), strongly indicated that besides petroleum hydrocarbons, plant wastes maybe available as possible sources of energy for resident soil bacteria at the China oil refineries site (Fig. 5, Supplementary Fig. S2, Supplementary Table S3, Supplementary Table S4). Bacteria are known to extracellularly depolymerize methylglucuronoxylan, a polysaccharide made of xylose that constitutes the hemicellulosic component of terrestrial plants ^51^ leading to the production of aldouronates and xylooligosaccharides. These compounds are taken up and normally converted intracellularly to fermentable xylose, leading to generation of energy along with ethanol. Alternatively, D-xylose can also be directly taken up from the environment. Also, two structural complexes for transport of peptides/oligopeptides (M00239 & M00439) were detected to be differentially abundant in the C samples along with bacterial proteasomes (M00342) (Fig. 5, Supplementary Fig. S2, Supplementary Table S3). This indicates that acquisition of environmental peptides and consequent proteasomal degradation of the same, may be the dominant mechanism for obtaining amino acids for assimilatory purposes in the C samples. Interestingly, the DesK-DesR two-component system, implicated in regulation of the *des* gene coding for a desaturase that helps control the saturation state of membrane lipids at low temperatures ^52^ was detected to be differentially abundant in the arctic samples (Fig. 5, Supplementary Fig. S2, Supplementary Table S3). Furthermore, the FitF-FitH two component system, responsible for insecticidal toxin regulation ^53^, was over-represented in the urban site of the Indian oil refinery samples (Fig. 5, Supplementary Fig. S2, Supplementary Table S3). This makes sense, since it has previously been shown that relatively higher amount of heat generation in cities compared to rural areas leads to sequestration of insects in urban areas ^54^. Sulfur assimilation in bacteria (M00616) was detected to be differentially abundant in OSC samples along with a number of modules dedicated to transfer of sulfur compounds (M00185, M00234, M00238, M00348, M00435-36) indicating a large genetic investment in scavenging and metabolism of sulfur compounds in this site (Fig. 5, Supplementary Fig. S2, Supplementary Table S3). Differential presence of a transport module for thiamine (M00191), which is required for assimilation of sulfonate compounds, furthers affirms this notion (Fig. 5, Supplementary Fig. S2, Supplementary Table S3). Additionally, differential detection of assimilatory nitrate reduction module (M00531) also indicates the availability and importance of nitrate ions in the sustenance of the OSC bacteriome (Fig. 5, Supplementary Fig. S2, Supplementary Table S3). Sulfate and nitrate ions are also important molecules in anaerobic respiration, and may therefore also play crucial roles in bacterial survival in the anaerobic regions of the OSC. Interestingly, reduction of nitrate has been reported to be closely linked to anaerobic degradation of benzene and concomitant growth ^55^, functional modules for both of which have been differentially detected in OSC samples (Fig. 5, Supplementary Fig. S2, Supplementary Table S3). Unlike other oil polluted sites, a large number of hydrocarbonoclastic modules differentially detected in the OSC samples (see previous section), whereas only two transport systems for small sugars (M00204, M00215) and no major polysaccharide metabolism and/or transport pathways were detected to be over-represented (Fig. 5, Supplementary Fig. S2, Supplementary Table S3). This not only indicates at the extreme habitat with highly restricted availability of carbon sources other than petroleum hydrocarbons in the OSC, it also explains the large clustering of differentially abundant hydrocarbon degradation pathways in the OSC samples and establishes petroleum hydrocarbons as the comprehensively dominant carbon assimilation/energy production source for this site. A number of functional modules related to methane metabolism, both methanogenic and methanotrophic, were detected to be over-represented for the OSTP samples. For example, methanogenesis (M00356) was over-represented in OSTPu, along with methane assimilation modules M00344-45 and M00608 detected to be differentially abundant in OSTPu and OSTPm respectively (Fig. 5, Supplementary Fig. S2, Supplementary Table S3). Oil sands tailings ponds are known to be important sources of methanogenesis and of methylotrophy ^17^, where deeper regions tend to be highly anaerobic. Additionally, modules for copper processing (M00762) and copper tolerance sensor (M00452) were detected in OSTPd (Fig. 5, Supplementary Fig. S2, Supplementary Table S3). This is important, since copper is an essential component of particulate methane monooxygenase (pMMO), and its availability can therefore determine the survivability of methanotrophs ^56^ along with the ratio of soluble and particulate MMO in the environment. A large number of modules dedicated to the biosynthesis of amino acids, vitamins and co-factors were detected to be over-represented in the taiga samples. This may be due to the poor availability of useful forms of amino acids and vitamins in the taiga environment. Presence of differentially abundant modules for sulfur containing amino acid biosynthesis (M00017, M00021) is supported by the detection of over-represented sulfur assimilation modules (M00176, M00595) which involve biosynthesis of cysteine and methionine as final/supplementary steps ^57,58^ (Fig. 5, Supplementary Fig. S2, Supplementary Table S3). Additionally, presence of alternative carbon sources as pectin and component sugars of other plant polysaccharides at the taiga sites can be inferred through the presence of differentially abundant functional modules for pectin degradation (M00081), and uptake and metabolism of other sugar and sugar derivatives as N-Acetylglucosamine, N, N’-Diacetylchitobiose, D-glucuronate, aldouronates, D-galactouronate (M00606, M00205, M00061, M00603, M00631) (Fig. 5, Supplementary Fig. S2, Supplementary Table S3). In the M samples, two component systems for starvation of phosphate (M00434), a limiting nutrient for mangroves ^59^ and metal tolerance (M00499) were detected as differentially abundant (Fig. 5, Supplementary Fig. S2, Supplementary Table S3). Genetic investment in metal tolerance should be important in M samples as mangroves in Brazil are routinely subjected to pollution from factory effluents ^42^. Furthermore, the clustering of differentially abundant central carbohydrate metabolism pathways (M00001-2, M00004, M00009, M00011) along with transport systems for fructose like sugars (M00273) in M samples, indicate availability of simple sugars as carbon sources besides hydrocarbons (Fig. 5, Supplementary Fig. S2, Supplementary Table S3). This is also supported by the differentially abundant module for synthesis of trehalose (M00565), a known carbohydrate energy storage compound and anti-desiccation agent ^60^, from glucose (Fig. 5, Supplementary Fig.S2, Supplementary Table S3). All functional modules for degradation of aromatic hydrocarbons were detected to be differentially abundant in OSC, DWH, taiga and OSTP (in that order) samples (Fig. 5, Supplementary Fig. S2, Supplementary Table S3). This is probably because these environments tend to be more extreme than other sites described in this study and coupled with oil pollution, the bacterial metabolic pathways in these environments have been further sculpted to rely greatly only on petroleum hydrocarbons for growth. Additionally, sulfate and nitrate utilization modules have been identified in most of these sites, which indicates an abundance of such ions in the environment and therefore use of the same for anaerobic alkane degradation, as previously described ^61^. Our results thus indicate that for all habitats, genetic composition of the bacteriome is representative of the immediate environment especially in terms of substrate usage, nutrient availability, energy metabolism, biosynthesis of compounds, and survival strategies including quorum sensing, chemotaxis, and stress response. Our findings reveal pathways differentially important in these oil polluted environments, especially those not related to hydrocarbon degradation and can therefore be used for differentiation between habitats of interest. Further empirical studies will however be required strengthen these observations and pinpoint functional biomarkers absolutely exclusive to oil polluted environments in specific biomes.

### Taxonomic biomarkers make important contributions to hydrocarbonoclastic and additional functional capacities in oil polluted environments

In order to identify taxonomic clades that may be differentially abundant in oil polluted sites used in the present study, taxonomic profiles generated through analysis of 16S rRNA data in mothur were examined using LEfSe. Additionally, to decipher functional associations of taxonomic clades, direct correlations between KOs and taxa were determined along with metagenomic contributions to hydrocarbonoclastic genes. Furthermore, bacterial co-occurrence and co-exclusion networks were deduced to understand important bacterial interactions in oil polluted sites. Our findings suggest that, taxonomic biomarkers inferred in our study contribute significantly to important functions in the oil polluted metabolic landscape and are often determined by their oil degradation capabilities. For example, biomarkers for DWH samples *HB2.32.21* and Alteromonadaceae (Fig. 4, Supplementary Table S2), were associated with over-represented KOs implicated in alginate biosynthesis (Supplementary Fig. S3). Moreover, a two component pathway involved in the regulation of alginate production (M00505) was also differentially abundant in DWH samples (Fig. 5, Supplementary Fig. S2, Supplementary Table S3). Interestingly, previous studies have shown that alginates provide increased mechanical stability to bacterial biofilms ^62^, and can therefore be instrumental in aiding anchorage or adhesion of DWH Alteromonadaceae. *HB.32.21* and Alteromonadaceae were found to be important contributors in hydrocarbonoclastic properties of the DWH bacteriome (Supplementary Fig. S4) and the former also exhibited strong associations with regulation of genes for aminobenzoate metabolism through *feaR* (see Results). Furthermore, another taxonomic biomarker identified for DWH samples, Colwelliaceae, was closely associated to the anaerobic C4-dicarboxylate transporter DcuB, which is responsible for transport of molecules as fumarate, succinate and malate ^63^. This is important, as it may help the facultatively aerobic Colwelliaceae to degrade alkanes anaerobically by addition of fumarates in marine sediments ^61^. Similarly, *Mycobacterium* was detected as a biomarker for C samples (Fig. 4, Supplementary Table S2) and correlated strongly with KOs implicated in degradation of hydrocarbons as naphthalene, benzoate and phthalate (Supplementary Fig. S3). *Mycobacterium* have previously been shown to harbor the ability to degrade a variety of aromatic hydrocarbons such as naphthalene, anthracene, phenanthrene, pyrene and so on ^32^. Phylum Cyanobacteria, a biomarker for C samples (Fig. 4, Supplementary Table S2), strongly correlated with differentially abundant photosynthetic proteins *cpeA* (K05376) and *psb28-2* (K08904) through an over-represented cyanobacterial order for C samples, Oscillatoriophycideae (Fig. 4, Supplementary Table S2). Additionally, KEGG modules for photosynthesis such as Photosystems I and II (M00163, M00161), cytochrome b6f complex (M00162) and NADP(H):quinone oxidoreductase for chloroplasts and cyanobacteria (M00145) were also found to be over-represented in C samples (Fig. 5, Supplementary Fig. S2, Supplementary Table S3). These observations indicated important, differential and extra-hydrocarbonoclastic contributions of Cyanobacteria in C samples. Furthermore, *Microbacterium,* which is known to be a stringent chemoorganotroph ^64^, was identified to be differentially abundant in A samples (Fig. 4, Supplementary Table S2). *Microbacterium* was found to share close associations with KOs involved in over-represented transport systems dedicated to acquisition of organic compounds such as cellobiose (M00206), alpha-glucosides (M00201), glutamate (M00227, M00233) and multiple sugars (M00207, M00216, M00221) (Supplementary Fig. S3), which can be used as possible sources of carbon and energy and also indicates availability of the same in the environment. Phylum Actinobacteria and class Actinobacteria, which were detected as biomarkers in the A samples, exhibited significant correlations with almost all differentially abundant KOs for A samples including hydrocarbon degrading genes as *pcaG,* phenol-2-monooxygenase, *bbsH*,and *pcaL* (data not shown). In OSC samples, over-represented taxa *Methylobacterium* was associated with genes involved in degradation of furfural and other hydrocarbons (Supplementary Fig. S3). The presence of the strongly aerobic *Methylobacterium* ^33^ once again reinforces the finding of ample availability of oxygen in the OSC. Interestingly, differentially abundant taxa Enterobacteriaceae and *Acinetobacter* were detected to be associated with a number of KOs implicated in stress response which included TCS KOs for aerobic/anaerobic survival as ArcB and NarP and transcriptional regulation of the *mar-sox-rob* regulon (Supplementary Fig. S3). The *mar-sox-rob* regulon has been reported in coordinating survival against various environmental stresses activated by inducers as paraquat, decanoate and intriguingly, salicylate ^65^, functional modules for which is differentially abundant in OSC samples (Fig. 5. Supplementary Fig. S2, Supplementary Table S3). Additionally, *Acinetobacter* was correlated with the stress response serine protease DegS and iron starvation Fe/S biogenesis protein NfuA (Supplementary Fig. S3). Thus, these biomarkers seem to contribute to important stress response pathways rather than hydrocarbon degrading capabilities. Additionally, over-represented taxa such as Oxalobacteraceae, *Cupriavidus, Brucellaceae*, and *Ochrobactrum* (Fig. 4, Supplementary Table S2) were found to differentially contribute to the abundance of a number of hydrocarbonoclastic genes (K00446, K00448-51, K03381) (Supplementary Fig. S4) in the OSC samples. In taiga samples, a number of detected biomarkers such as *Phenylobacterium*, Caulobacteraceae, Sphingomonadaceae, *Novosphingobium*, *Rhodococcus*, and Burkholderiaceae (Fig. 4, Supplementary Table S2) were found to contribute heavily but differently to the abundance of a plethora of hydrocarbonoclastic genes (Supplementary Fig. S4). Differentially abundant functional modules for the assimilation of sulphate, transformation of thiosulphate to sulphate and regulation of the SOX complex responsible for thiosulphate transformation (M00176, M00595, M00523) underline the preferential sulphur usage in this site (Fig. 5, Supplementary Fig. S2, Supplementary Table S3). This is well supported by the identification of *Bradyrhizobium*, *Caulobacter*, and *Burkholderia* as biomarkers (Fig. 4, Supplementary Table S2), all which are known to be involved in sulfur metabolism ^66,67^ and house homologous genes for the same. Interestingly, a number of biomarkers identified here for the taiga samples such as *Phenylobacterium*, Sphingomonadaceae, *Novosphingobium* and *Rhodococcus* were detected as “habitat specialists” in oil contaminated taiga samples by Yang et al. ^68^. Similar to the results of An et al. ^17^, we encountered a significantly high proportion of anaerobic taxa in the OSTP samples, among which Anaerolinaceae, Syntrophaceae, Desulfobulbaceae, Peptococcaceae, Geobacteraceae, Syntrophorhabdaceae and the thermophilic Caldiserica ^69^ were detected as biomarkers (Fig. 4, Supplementary Table S2). Detected taxonomic biomarkers such as Anaerolinaceae and Comamonadaceae (Fig. 4, Supplementary Table S2) were found to make significant contributions to the abundance of hydrocarbon degradation genes (Supplementary Fig. S4) in OSTP samples. Other identified biomarkers such as Geobacteraceae and *Thauera* (Fig. 4, Supplementary Table S2) are well known anaerobic hydrocarbon degraders ^70,71^. Additionally, another detected biomarker Nitrospirales (Fig. 4, Supplementary Table S2), which is involved in nitrification ^72^ may contribute to nitrification, for which over-represented module ammonia => nitrite transformation (M00528) was identified in OSTP samples (Fig. 5, Supplementary Fig. S2, Supplementary Table S3). Biomarkers of sulfate reducing bacteria such as Desulfuromonadales and Desulfobulbaceae (Fig. 4, Supplementary Table S2), which is a known mesophilic/psychrophilic sulfate reducer ^73^ may be involved in important sulfur metabolism pathways known to be important in OSTPs ^74^. Interestingly, obligate anaerobes Anaerolinaceae have previously been associated with sulfate reducing conditions in the OSTPs ^75^. Lastly, major contributions for hydrocarbonoclastic capabilities in OSTP samples was also observed from biomarkers *Pseudomonas* (K00446, K00448, K00449, K00496, K03381) and Rhodocyclaceae (K04100-01) (Fig. 4, Supplementary Table S2, Supplementary Fig. S4), furthering the hydrocarbon degradation capabilities of OSTPs.

We also investigated significant bacterial associations in oil polluted sites to decipher important co-occurrence and co-exclusions. Our results showed that greater co-occurrence exists between phylotypes sharing an evolutionary lineage while more co-exclusions were observed between phylotypes from different ancestries. This observation has also been previously reported in microbial correlation studies in the environment ^76,77^. Interestingly, not a large proportion of taxonomic biomarkers were observed to be represented in these significant correlations. This can possibly happen due to separation of niches due to various environmental and even temporal factors. For example, in the bacterial association network for DWH samples, biomarker Colwelliaceae was detected to participate in a significantly positive relationship with Desulfobulbaceae, a strictly anaerobic sulfate utilizing bacterial family (Fig. 6). The existence of this kind of a relationship, based on degradation of recalcitrant hydrocarbons, was inferred upon by the original authors too ^47^. Strikingly however, the most abundant and robust hydrocarbon degrader i.e. *HB2.32.21* (Supplementary Fig. S4) was not detected to be involved in any significant associations. This observation can be explained by a possible individual capacity of survival for *HB2.32.21* due to its hydrocarbonoclastic capacities without extensive interactions with other resident bacteria, therefore occupying a separate niche in the oil polluted marine sediment site. Thus, significant correlations (both positive and negative) may be driven by factors other than only oil pollution in oil contaminated sites with apparently benign taxa being involved in such interactions. This indicates that biomarkers and correlation networks must be studied in tandem to deduce meaningful conclusions. Even though empirical evidence in support of the natural presence of most microbial association networks is lacking, our study shows that they may be important to holistic interpretation of results as well as being a good starting point for further investigations.

Our results therefore show that, detected biomarkers may contribute differently to strictly hydrocarbonoclastic properties when compared across sites, but their close association with a majority of differentially abundant KOs and as an extension a number of over-represented functional pathways for each site underlines their significance in these oil contaminated sites. We find that although many of the taxonomic biomarkers contribute to hydrocarbonoclastic capacities, some do not and can therefore contribute to other possibly important functions. These observations not only elucidate important taxa contributing functions more specific and essential to each site, but also shows that niches related to functions other than hydrocarbon degradation may significantly influence bacteriome structure in oil polluted sites, possibly more in sites with understandably lower degrees of contamination. This indicates clearly that while hydrocarbonoclastic capabilities may be a driving force for continued survival in these sites, other immediate factors including availability of different organic and inorganic compounds and environmental stress play heavily on the evolution of the bacteriome. Thus, we see that a combination of oil degradation capabilities and environmental factors shape the landscape for bacterial petroleum degradation. As an extension, it therefore becomes imperative to examine oil bioremediation processes, especially aimed at empirical identification of biomarkers, in totality with due comparison to similar studies and not in isolation as it may lead to misleading conclusions. This is well illustrated in some previous studies that have focused on predicting microbial markers or proxies for oil pollution in certain environments ^42,43^. In the study on mangrove oil pollution and detection of microbial proxies by dos Santos et al. ^43^, *Marinobacter*, belonging to family Alteromonadaceae, was identified as a possible biomarker for oil pollution in mangroves. However, in our study when compared to other sites, Alteromonadaceae was detected to be differentially abundant in the DWH samples and *Marinobacter* was not identified as over-represented in any of the oil polluted sites (Fig. 4, Supplementary Table S2).

## Conclusion

In summary, our study showed that significant taxonomic and functional differences exist between geographically and/or spatially isolated oil polluted sites and that oil pollution is not the sole driving factor in determination of the bacteriome at these sites, even if maybe the most predominant one. In our study we have successfully detected functions that are important and contribute significantly to the sustenance of bacteriomes at these oil polluted sites. Additionally, we have detected taxa that are differentially abundant and also contribute to many of these functions. Furthermore, we have also successfully shown that several of these important taxonomic clades and functional modules are often involved in extra-hydrocarbonoclastic activities, thus underlining the importance of these apparently peripheral niches related to endemic environmental responses such as variable resource utilization, alternate respiration and stress response in the survival of oil contaminated ecosystems. In the process, we therefore identified robust taxonomic and functional biomarkers, that are representative of an entire oil polluted environment and not only its hydrocarbonoclastic capabilities. These biomarkers can be implemented in monitoring of bacterial remediation processes as well as for distinguishing one of the 12 oil polluted habitats. Our results show that some parallels exist in the functional composition of oil polluted environments, mainly regarding adaptations to aerobic and anaerobic lifestyles depending on availability of limited resources compatible with sustenance of anaerobic growth (e.g. sulfate/nitrate related metabolism), but differences between them pertaining to transport of substrates, biosynthesis of biomolecules, response to various stress, degradation of diverse compounds and so on are significant and together with differences identified in taxonomic profiles and hydrocarbonoclastic capabilities enable us to truly differentiate between them. However, further studies are required which may involve simultaneous 16S rRNA based phylogenetic survey, metagenomic and metatranscriptomic investigations of oil polluted environments followed by empirical experimentations on the same to confirm robust and significantly differential taxonomic and functional biomarkers which may be used for monitoring. With the current sequencing technologies, bioinformatics tools and increasing data from other studies, it is highly possible that characteristic degradation profiles and subsequent remediation strategies for different environments across the world will be inferred fairly soon with unprecedented confidence. To our knowledge, this is the first population genomics study carried out on petroleum hydrocarbon polluted habitats. Our study presents novel understanding of oil contaminated habitats by showing how and in what manner petroleum hydrocarbons fashion the metagenomic fabric along with the evident effect of endemic factors characteristic of geographically separated diverse petroleum hydrocarbon contaminated environments, while interpreting the effects of both on the resident microbiome. This study therefore provides a foundation for future investigations into the microbial biodiversity and habitat function in oil polluted sites.

## Acknowledgements

The authors acknowledge the financial support provided for the research presented above from the Department of Biotechnology, Government of India vide Sanction no. BT/306/NE/TBP/2012 dated December 6, 2012, under the DBT-Twinning scheme. A.M. was supported by the CSIR/UGC-NET Fellowship from the University Grants Commission, Government of India (201112-NETJRF-10217-100). We thank the Bioinformatics Resources and Applications Facility (BRAF), CDAC, Pune and the Centre for High Performance Computing for Modern Biology, University of Calcutta for granting access to supercomputing facilities along with their unconditional and continuous help. We would also like to thank the ICZMP, World Bank and Department of Biochemistry, University of Calcutta for allowing us to use the Pyrosequencing facility. Lastly, we would like to acknowledge the DST-Purse, Department of Biotechnology, University of Calcutta and the University of Calcutta for providing the necessary infrastructure for implementation of the research work furnished above.

## Author contributions

D.C. and A.K.S. managed the project. A.M., D.C. and A.K.S. conceptualized and designed the experiments. B.C., J.L., A.K.S. and A.K.M. designed and conducted sampling for oil contaminated soil. A.M., B.C., J.L., P.B. and M.B. were involved in designing the sequencing strategy and conducting the same. A.M. designed the bioinformatic analysis strategy and conducted the same with assistance from A.P. A.M. and D.C. performed the data analyses. A.M, D.C. and A.K.S. prepared the manuscript. All authors reviewed the manuscript.

## References

1 Dean-Ross, D., Moody, J. & Cerniglia, C. E. Utilization of mixtures of polycyclic aromatic hydrocarbons by bacteria isolated from contaminated sediment. FEMS microbiology ecology 41, 1–7, doi:10.1111/j.1574-6941.2002.tb00960.x (2002).

2 Molina, M., Araujo, R. & Hodson, R. E. Cross-induction of pyrene and phenanthrene in a Mycobacterium sp. isolated from polycyclic aromatic hydrocarbon contaminated river sediments. Canadian journal of microbiology 45, 520–529 (1999).

3 Stringfellow, W. T. & Aitken, M. D. Competitive metabolism of naphthalene, methylnaphthalenes, and fluorene by phenanthrene-degrading pseudomonads. Applied and environmental microbiology 61, 357–362 (1995).

4 Bakken, L. R. Culturable and non-culturable bacteria in soil. J.D. van Elsas, J.T. Trevor, E.M.H. Wellington (Eds.), Modern soil microbiology, Marcel Dekker, New York 47–61 (1997).

5 Gevers, D. et al. The Human Microbiome Project: a community resource for the healthy human microbiome. PLoS biology 10, e1001377, doi:10.1371/journal.pbio.1001377 (2012).

6 Segata, N. et al. Computational meta'omics for microbial community studies. Molecular systems biology 9, 666, doi:10.1038/msb.2013.22 (2013).

7 Friedman, J. & Alm, E. J. Inferring correlation networks from genomic survey data. PLoS computational biology 8, e1002687, doi:10.1371/journal.pcbi.1002687 (2012).

8 Segata, N. et al. Metagenomic biomarker discovery and explanation. Genome biology 12, R60, doi:10.1186/gb-2011-12-6-r60 (2011).

9 Langille, M. G. et al. Predictive functional profiling of microbial communities using 16S rRNA marker gene sequences. Nature biotechnology 31, 814–821, doi:10.1038/nbt.2676 (2013).

10 Weisburg, W. G., Barns, S. M., Pelletier, D. A. & Lane, D. J. 16S ribosomal DNA amplification for phylogenetic study. Journal of bacteriology 173, 697–703 (1991).

11 Lofgren, J. L. et al. Lack of commensal flora in Helicobacter pylori-infected INS-GAS mice reduces gastritis and delays intraepithelial neoplasia. Gastroenterology 140, 210–220, doi:10.1053/j.gastro.2010.09.048 (2011).

12 Andrews, S. FastQC: a quality control tool for high throughput sequence data. Available online at: http://www.bioinformatics.babraham.ac.uk/projects/fastqc (2010).

13 Schloss, P. D. et al. Introducing mothur: open-source, platform-independent, community-supported software for describing and comparing microbial communities. Applied and environmental microbiology 75, 7537–7541, doi:10.1128/AEM.01541-09 (2009).

14 Edgar, R. C., Haas, B. J., Clemente, J. C., Quince, C. & Knight, R. UCHIME improves sensitivity and speed of chimera detection. Bioinformatics 27, 2194–2200, doi:10.1093/bioinformatics/btr381 (2011).

15 DeSantis, T. Z. et al. Greengenes, a chimera-checked 16S rRNA gene database and workbench compatible with ARB. Applied and environmental microbiology 72, 5069–5072, doi:10.1128/AEM.03006-05 (2006).

16 Asnicar, F., Weingart, G., Tickle, T. L., Huttenhower, C. & Segata, N. Compact graphical representation of phylogenetic data and metadata with GraPhlAn. PeerJ 3, e1029, doi:10.7717/peerj.1029 (2015).

17 An, D. et al. Metagenomics of hydrocarbon resource environments indicates aerobic taxa and genes to be unexpectedly common. Environmental science & technology 47, 10708–10717, doi:10.1021/es4020184 (2013).

18 Yang, S., Wen, X., Zhao, L., Shi, Y. & Jin, H. Crude oil treatment leads to shift of bacterial communities in soils from the deep active layer and upper permafrost along the China-Russia Crude Oil Pipeline route. PloS one 9, e96552, doi:10.1371/journal.pone.0096552 (2014).

19 Bray, J. R. & Curtis, J. T. An ordination of the upland forest communities of southern Wisconsin. Ecol Monographs 27, 325–349 (1957).

20 Hammer, Ø., Harper, D. A. T. & Ryan, P. D. PAST: Paleontological Statistics Software Package for Education and Data Analysis. Palaeontologia Electronica 4, 9pp (2001).

21 Markowitz, V. M. et al. IMG: the Integrated Microbial Genomes database and comparative analysis system. Nucleic acids research 40, D115–122, doi:10.1093/nar/gkr1044 (2012).

22 Kanehisa, M., Goto, S., Furumichi, M., Tanabe, M. & Hirakawa, M. KEGG for representation and analysis of molecular networks involving diseases and drugs. Nucleic acids research 38, D355–360, doi:10.1093/nar/gkp896 (2010).

23 Wickham, H. ggplot2: Elegant Graphics for Data Analysis. Springer-Verlag New York (2009).

24 Abubucker, S. et al. Metabolic reconstruction for metagenomic data and its application to the human microbiome. PLoS computational biology 8, e1002358, doi:10.1371/journal.pcbi.1002358 (2012).

25 Ye, Y. & Doak, T. G. A parsimony approach to biological pathway reconstruction/inference for genomes and metagenomes. PLoS computational biology 5, e1000465, doi:10.1371/journal.pcbi.1000465 (2009).

26 Goecks, J., Nekrutenko, A., Taylor, J. & Galaxy, T. Galaxy: a comprehensive approach for supporting accessible, reproducible, and transparent computational research in the life sciences. Genome biology 11, R86, doi:10.1186/gb-2010-11-8-r86 (2010).

27 Revelle, W. psych: Procedures for Personality and Psychological Research. (2016).

28 Shannon, P. et al. Cytoscape: a software environment for integrated models of biomolecular interaction networks. Genome research 13, 2498–2504, doi:10.1101/gr.1239303 (2003).

29 Joshi, M. N. et al. Metagenomic approach for understanding microbial population from petroleum muck. Genome announcements 2, doi:10.1128/genomeA.00533-14 (2014).

30 Yergeau, E., Sanschagrin, S., Beaumier, D. & Greer, C. W. Metagenomic analysis of the bioremediation of diesel-contaminated Canadian high arctic soils. PloS one 7, e30058, doi:10.1371/journal.pone.0030058 (2012).

31 Segata, N. et al. Composition of the adult digestive tract bacterial microbiome based on seven mouth surfaces, tonsils, throat and stool samples. Genome biology 13, R42, doi:10.1186/gb-2012-13-6-r42 (2012).

32 Kim, S. J., Kweon, O. & Cerniglia, C. E. Degradation of Polycyclic Aromatic Hydrocarbons by Mycobacterium Strains. Handbook of Hydrocarbon and Lipid Microbiology, 1865–1879.

33 Green, P. N. Methylobacterium. The Prokaryotes: Proteobacteria: Alpha and Beta Subclasses 5, 257–265, doi:10.1007/0-387-30745-1_14 (2006).

34 Jurkowski, A., Reid, A. H. & Labov, J. B. Metagenomics: a call for bringing a new science into the classroom (while it’s still new). CBE life sciences education 6, 260–265 (2007).

35 Guo, J. et al. Dissecting microbial community structure and methane-producing pathways of a full-scale anaerobic reactor digesting activated sludge from wastewater treatment by metagenomic sequencing. Microbial cell factories 14, 33 (2015).

36 He, Y., Feng, X., Fang, J., Zhang, Y. & Xiao, X. Metagenome and Metatranscriptome Revealed a Highly Active and Intensive Sulfur Cycle in an Oil-Immersed Hydrothermal Chimney in Guaymas Basin. Frontiers in microbiology 6, 1236 (2015).

37 Kuppusamy, S. et al. Pyrosequencing analysis of bacterial diversity in soils contaminated long-term with PAHs and heavy metals: Implications to bioremediation. Journal of hazardous materials 317, 169–179 (2016).

38 Cerqueira, T. et al. Microbial diversity in deep-sea sediments from the Menez Gwen hydrothermal vent system of the Mid-Atlantic Ridge. Marine genomics 24 Pt 3, 343–355, doi:10.1016/j.margen.2015.09.001 (2015).

39 Tytgat, B. et al. Bacterial diversity assessment in Antarctic terrestrial and aquatic microbial mats: a comparison between bidirectional pyrosequencing and cultivation. PloS one 9, e97564, doi:10.1371/journal.pone.0097564 (2014).

40 Barton, H. A. et al. Microbial diversity in a Venezuelan orthoquartzite cave is dominated by the Chloroflexi (Class Ktedonobacterales) and Thaumarchaeota Group I.1c. Frontiers in microbiology 5, 615, doi:10.3389/fmicb.2014.00615 (2014).

41 Mukherjee, A. & Chattopadhyay, D. Exploring environmental systems and processes through next-generation sequencing technologies: insights into microbial response to petroleum contamination in key environments. The Nucleus, doi:10.1007/s13237-016-0190-3 (2016).

42 Andreote, F. D. et al. The microbiome of Brazilian mangrove sediments as revealed by metagenomics. PloS one 7, e38600, doi:10.1371/journal.pone.0038600 (2012).

43 dos Santos, H. F. et al. Mangrove bacterial diversity and the impact of oil contamination revealed by pyrosequencing: bacterial proxies for oil pollution. PloS one 6, e16943, doi:10.1371/journal.pone.0016943 (2011).

44 Hernandez-Raquet, G. et al. Molecular diversity studies of bacterial communities of oil polluted microbial mats from the Etang de Berre (France). FEMS microbiology ecology 58, 550–562, doi:10.1111/j.1574-6941.2006.00187.x (2006).

45 Kane, S. R. et al. Whole-genome analysis of the methyl tert-butyl ether-degrading beta-proteobacterium Methylibium petroleiphilum PM1. Journal of bacteriology 189, 1931–1945, doi:10.1128/JB.01259-06 (2007).

46 Rosenberg, E. The Family Chitinophagaceae. The Prokaryotes: Other Major Lineages of Bacteria and The Archaea, 493–495, doi:10.1007/978-3-642-38954-2_137 (2014).

47 Mason, O. U. et al. Metagenomics reveals sediment microbial community response to Deepwater Horizon oil spill. The ISME journal 8, 1464–1475, doi:10.1038/ismej.2013.254 (2014).

48 Jindrova, E., Chocova, M., Demnerova, K. & Brenner, V. Bacterial aerobic degradation of benzene, toluene, ethylbenzene and xylene. Folia microbiologica 47, 83–93 (2002).

49 Pelletier, D. A. & Harwood, C. S. 2-Hydroxycyclohexanecarboxyl coenzyme A dehydrogenase, an enzyme characteristic of the anaerobic benzoate degradation pathway used by Rhodopseudomonas palustris. Journal of bacteriology 182, 2753–2760 (2000).

50 Wood, J. M. Osmosensing by bacteria: signals and membrane-based sensors. Microbiology and molecular biology reviews: MMBR 63, 230–262 (1999).

51 Chow, V., Nong, G. & Preston, J. F. Structure, function, and regulation of the aldouronate utilization gene cluster from Paenibacillus sp. strain JDR-2. Journal of bacteriology 189, 8863–8870, doi:10.1128/JB.01141-07 (2007).

52 Mansilla, M. C., Cybulski, L. E., Albanesi, D. & de Mendoza, D. Control of membrane lipid fluidity by molecular thermosensors. Journal of bacteriology 186, 6681–6688, doi:10.1128/JB.186.20.6681-6688.2004 (2004).

53 Kupferschmied, P., Pechy-Tarr, M., Imperiali, N., Maurhofer, M. & Keel, C. Domain shuffling in a sensor protein contributed to the evolution of insect pathogenicity in plant-beneficial Pseudomonas protegens. PLoS pathogens 10, e1003964, doi:10.1371/journal.ppat.1003964 (2014).

54 Meineke, E. K., Dunn, R. R., Sexton, J. O. & Frank, S. D. Urban warming drives insect pest abundance on street trees. PloS one 8, e59687, doi:10.1371/journal.pone.0059687 (2013).

55 Burland, S. M. & Edwards, E. A. Anaerobic benzene biodegradation linked to nitrate reduction. Applied and environmental microbiology 65, 529–533 (1999).

56 Lieberman, R. L. & Rosenzweig, A. C. Biological methane oxidation: regulation, biochemistry, and active site structure of particulate methane monooxygenase. Critical reviews in biochemistry and molecular biology 39, 147–164, doi:10.1080/10409230490475507 (2004).

57 Wei, J. et al. Cysteine biosynthetic enzymes are the pieces of a metabolic energy pump. Biochemistry 41, 8493–8498 (2002).

58 Sekowska, A., Kung, H. F. & Danchin, A. Sulfur metabolism in Escherichia coli and related bacteria: facts and fiction. Journal of molecular microbiology and biotechnology 2, 145–177 (2000).

59 Chakraborty, A. et al. Changing bacterial profile of Sundarbans, the world heritage mangrove: impact of anthropogenic interventions. World journal of microbiology & biotechnology 31, 593–610, doi:10.1007/s11274-015-1814-5 (2015).

60 Elbein, A. D., Pan, Y. T., Pastuszak, I. & Carroll, D. New insights on trehalose: a multifunctional molecule. Glycobiology 13, 17R–27R, doi:10.1093/glycob/cwg047 (2003).

61 Rojo, F. Degradation of alkanes by bacteria. Environmental microbiology 11, 2477–2490, doi:10.1111/j.1462-2920.2009.01948.x (2009).

62 Garrett, T. R., Bhakoo, M. & Zhang, Z. B. Bacterial adhesion and biofilms on surfaces. Prog Nat Sci 18, 1049–1056, doi:10.1016/j.pnsc.2008.04.001 (2008).

63 Ullmann, R., Gross, R., Simon, J., Unden, G. & Kroger, A. Transport of C(4)-dicarboxylates in Wolinella succinogenes. Journal of bacteriology 182, 5757–5764 (2000).

64 Suzuki, K. I. & Hamada, M. Microbacterium. Bergey’s Manual of Systematics of Archaea and Bacteria, 1–52, doi:10.1002/9781118960608.gbm00104 (2015).

65 Chubiz, L. M., Glekas, G. D. & Rao, C. V. Transcriptional cross talk within the mar-sox-rob regulon in Escherichia coli is limited to the rob and marRAB operons. Journal of bacteriology 194, 4867–4875, doi:10.1128/JB.00680-12 (2012).

66 Elsen, S., Swem, L. R., Swem, D. L. & Bauer, C. E. RegB/RegA, a highly conserved redox-responding global two-component regulatory system. Microbiology and molecular biology reviews: MMBR 68, 263–279, doi:10.1128/MMBR.68.2.263-279.2004 (2004).

67 Lochowska, A. et al. Regulation of sulfur assimilation pathways in Burkholderia cenocepacia through control of genes by the SsuR transcription factor. Journal of bacteriology 193, 1843–1853, doi:10.1128/JB.00483-10 (2011).

68 Yang, S. et al. Hydrocarbon degraders establish at the costs of microbial richness, abundance and keystone taxa after crude oil contamination in permafrost environments. Scientific Reports 6, 37473, doi:10.1038/srep37473 (2016).

69 Mori, K., Yamaguchi, K., Sakiyama, Y., Urabe, T. & Suzuki, K. Caldisericum exile gen. nov., sp. nov., an anaerobic, thermophilic, filamentous bacterium of a novel bacterial phylum, Caldiserica phyl. nov., originally called the candidate phylum OP5, and description of Caldisericaceae fam. nov., Caldisericales ord. nov. and Caldisericia classis nov. International journal of systematic and evolutionary microbiology 59, 2894–2898, doi:10.1099/ijs.0.010033-0 (2009).

70 Childers, S. E., Ciufo, S. & Lovley, D. R. Geobacter metallireducens accesses insoluble Fe(III) oxide by chemotaxis. Nature 416, 767–769, doi:10.1038/416767a (2002).

71 Macy, J. M. et al. Thauera selenatis gen. nov., sp. nov., a member of the beta subclass of Proteobacteria with a novel type of anaerobic respiration. International journal of systematic bacteriology 43, 135–142, doi:10.1099/00207713-43-1-135 (1993).

72 Daims, H. et al. Complete nitrification by Nitrospira bacteria. Nature 528, 504–509, doi:10.1038/nature16461 (2015).

73 Kuever, J. The Family Desulfobulbaceae. The Prokaryotes: Deltaproteobacteria and Epsilonproteobacteria, 75–86, doi:10.1007/978-3-642-39044-9_267 (2014).

74 Warren, L. A., Kendra, K. E., Brady, A. L. & Slater, G. F. Sulfur Biogeochemistry of an Oil Sands Composite Tailings Deposit. Frontiers in microbiology 6, 1533, doi:10.3389/fmicb.2015.01533 (2015).

75 Penner, T. J. & Foght, J. M. Mature fine tailings from oil sands processing harbour diverse methanogenic communities. Canadian journal of microbiology 56, 459–470, doi:10.1139/w10-029 (2010).

76 Xu, Z., Hansen, M. A., Hansen, L. H., Jacquiod, S. & Sorensen, S. J. Bioinformatic approaches reveal metagenomic characterization of soil microbial community. PloS one 9, e93445, doi:10.1371/journal.pone.0093445 (2014).

77 Barret, M. et al. Emergence shapes the structure of the seed microbiota. Applied and environmental microbiology 81, 1257–1266, doi:10.1128/AEM.03722-14 (2015).

78 Bell, T. H. et al. Predictable bacterial composition and hydrocarbon degradation in Arctic soils following diesel and nutrient disturbance. The ISME journal 7, 1200–1210, doi:10.1038/ismej.2013.1 (2013).

79 Sun, W. et al. Microbial communities inhabiting oil-contaminated soils from two major oilfields in Northern China: Implications for active petroleum-degrading capacity. Journal of microbiology 53, 371–378, doi:10.1007/s12275-015-5023-6 (2015).

